# iPSC-Derived Chondroprogenitors as a Promising Cell Source for Cartilage Engineering: A Comparison with MSCs in Unmodified and Peptide-Functionalized Alginate Hydrogels

**DOI:** 10.64898/2026.09.08.750074

**Authors:** Maria Laura Vieri, Andrew R. Thomson, Matthew J. Dalby, Nikola Kolundzic, Cecilia Ansalone, Bart Dietrich, Penelope M. Tsimbouri, Tyler Shaw, Carmen Huesa, Dave J. Adams, Carl S. Goodyear

## Abstract

Articular cartilage degeneration is a hallmark of degenerative joint diseases, yet its limited regenerative capacity poses significant challenges for tissue engineering. While mesenchymal stem cells (MSCs) are the most commonly employed cell type in cartilage tissue engineering, they exhibit donor variability, restricted expansion capacity, and tendencies toward fibrocartilage formation and hypertrophic differentiation. Human induced pluripotent stem cell-derived chondroprogenitors (iCPs) represent a promising alternative, offering scalable production of developmentally relevant cells with intrinsic chondrogenic commitment. However, their performance within three-dimensional biomaterial scaffolds remains largely unexplored. Here, we compared chondrogenic differentiation of iCPs and MSCs within alginate hydrogels of varying stiffness (0.37–4.55 kPa) exhibiting physiologically relevant stress relaxation properties. Intermediate stiffness (2% alginate, ∼2.17 kPa) optimally supported chondrogenesis for both cell types. While MSCs differentiated as single cells, iCPs spontaneously self-organized into cartilaginous aggregates without requiring a separate pellet pre-culture step, showing significantly higher hyaline indices and reduced COL10 expression, despite initial low viability in hydrogels. To further enhance chondrogenesis, we functionalized 2% alginate gels with RGD and HAVDI peptides mimicking integrin- and cadherin-mediated signaling. HAVDI/RGD functionalization significantly enhanced hyaline cartilage marker expression in both cell types, with iCPs exhibiting superior matrix composition characterized by elevated aggrecan and SOX9 expression and reduced COL10 and MMP13 compared to MSCs. These findings establish iCPs as a promising cell source for cartilage tissue engineering and disease modeling, particularly within biomaterials integrating mechanical and bioactive cues that recapitulate the native cartilage microenvironment.

**HIGHLIGHTS:**

- First direct comparison of iPSCs-derived chondroprogenitors (iCPs) vs MSCs in peptide-functionalized 3D hydrogels.
- Soft, fast-relaxing 2% (w/v) alginate gels support chondrogenesis of both cell types: iCPs form cartilaginous aggregates, MSCs differentiate as single cells.
- HAVDI/RGD peptides enhance hyaline cartilage marker expression and ECM deposition in both cell types and improve iCPs cellular organization.
- iCPs-derived cartilage in engineered alginate hydrogel exhibits more native-like matrix composition and organization compared to MSCs.

**GRAPHICAL ABSTRACT:** 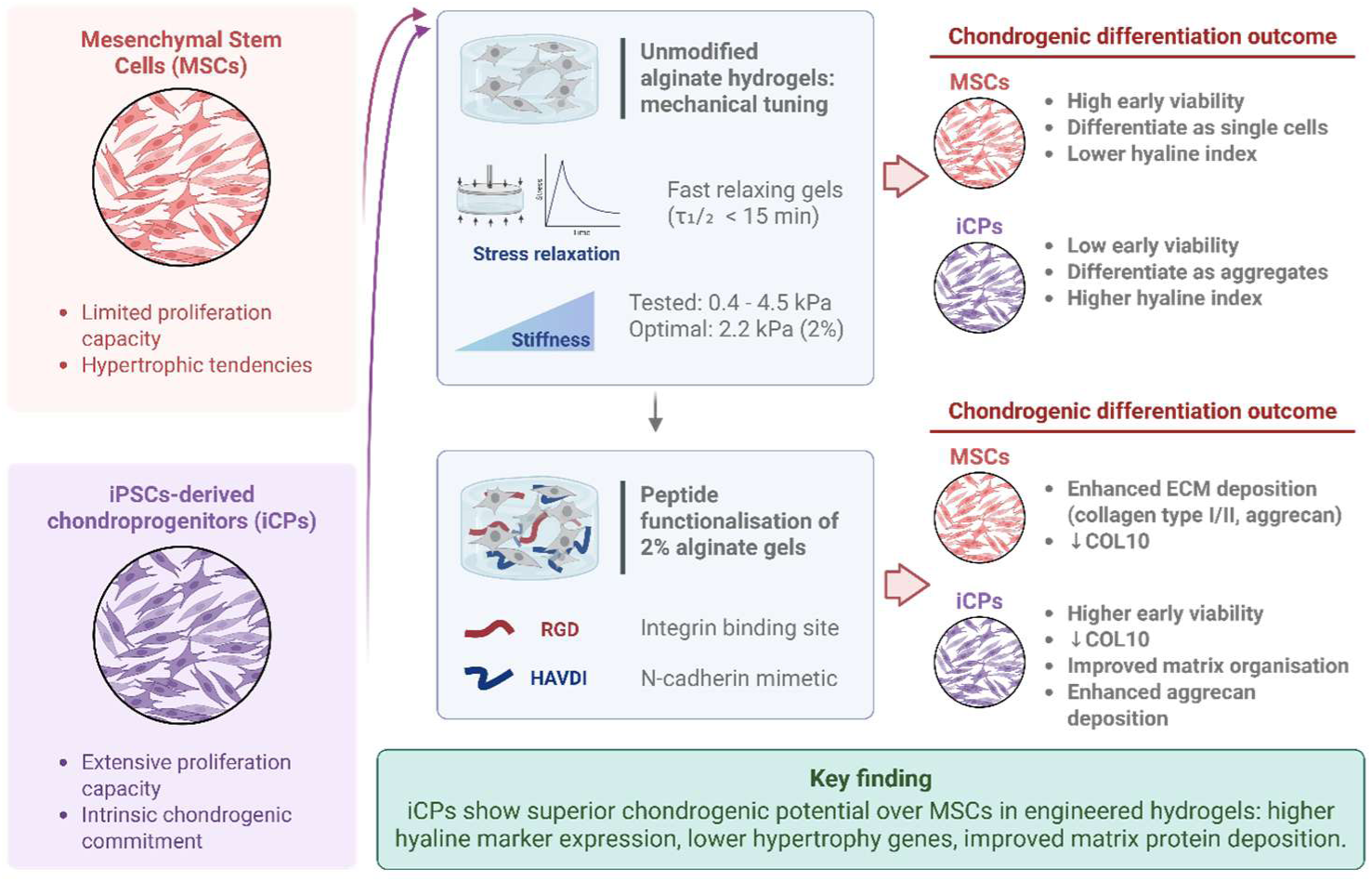

## 1. INTRODUCTION

Articular cartilage degeneration is the hallmark of degenerative joint diseases, including osteoarthritis (OA), rheumatoid arthritis (RA), and spondyloarthropathies (SpA), which represent one of the leading causes of disability globally, placing a substantial burden on healthcare systems and economies [1]. Given its avascular nature, low cellularity, and limited regenerative ability, articular cartilage represents a challenging tissue to regenerate and to study in vitro through tissue engineering approaches [2–4]. Cartilage tissue engineering has advanced significantly in the past decade, providing innovative biomaterial scaffolds that support stem cell differentiation into cartilage [5–7]. However, identifying the optimal cell source and establishing robust protocols for stem cell isolation, expansion, and differentiation remain crucial for achieving effective chondrogenesis [5,8,9].

Mesenchymal stem cells (MSCs) are one of the most used cells in cartilage tissue engineering, due to their relative abundance and remarkable capacity to differentiate into various cell types, including chondrocytes [10]. Typically sourced from bone marrow and adipose tissue [10–12], MSCs have been widely used in promoting chondrogenesis in both pellet cultures and three-dimensional (3D) scaffolds [13–15]. Despite their utility, MSCs face limitations such as variability due to donor characteristics, tissue source, isolation methods, and in vitro culture conditions [15–18]. These sources of variability, along with the limited expansion capacity of MSCs [19,20], constrain their reproducibility and scalability for cartilage tissue engineering. Additionally, MSC-derived tissues tend to display fibrocartilage-like characteristics and undergo hypertrophy [13,21–23] rather than form hyaline cartilage, limiting their suitability for modelling native cartilage. These limitations suggest that MSC functions and responses do not fully recapitulate developmental chondrogenesis, highlighting the need for more rigorous methodologies to mimic embryonic cartilage formation more closely [22,24].

Chondroprogenitors (CPs), located within articular cartilage, have shown considerable potential for cartilage regeneration and tissue engineering due to their intrinsic commitment towards the chondrogenic lineage and lower expression of hypertrophy markers [25–28]. Human CPs are characterized by CD146 and CD166 expression and absence of CD34 [25], and exhibit increased chondrogenic activity compared to human MSCs [26]. Consequently, CPs may represent a more suitable cell source than adult dedifferentiated chondrocytes or MSCs [29,30]. However, identifying a practical and abundant source of expandable CPs for widespread use remains challenging due to gaps in their biological profile, variability in isolation and expansion methods, and their rarity as a cell population [26,27,31].

Human induced pluripotent stem cells (iPSCs) represent a potential source of CPs, enabling their generation through the recapitulation of embryonic developmental pathways [32,33]. Generated through the reprogramming of somatic cells, iPSCs offer a more controlled and potentially more reproducible alternative for cartilage tissue engineering and regenerative therapies because of their pluripotency, extensive proliferation capacity and ability to generate lineage-specific progenitors under defined culture conditions [14,34], while also avoiding some of the ethical concerns associated with embryonic stem cells (ESCs) [35]. Additionally, they can be genetically engineered, enabling controlled disease modelling of musculoskeletal disorders OA [36–38]. As iPSCs provide a scalable source of developmentally derived CPs, iPSCs-derived chondroprogenitors (iCPs) represent a defined cell population for cartilage tissue engineering and musculoskeletal disease modelling [29,33,39–41].

In this study, we assessed human iCPs as a cell source for cartilage tissue engineering within 3D biomaterial scaffolds and compare their performance to human MSCs. While iCPs have recently been shown to outperform MSCs in pellet cultures [42], their behavior in 3D biomaterial scaffold environments remains largely unexplored. Alginate hydrogels were selected as the scaffold platform due to their biocompatibility, cost-effectiveness, rapid gelation, tunable mechanical properties, and capacity for chemical functionalization [43,44]. We then identified the optimal mechanical properties of alginate gels for iCPs chondrogenesis, by analyzing gel viscoelasticity and stress relaxation [45–48] using rheological methods. Developmentally relevant peptides were subsequently incorporated to modulate stem cell chondrogenesis, building on previous observations in MSCs systems [49–54]. Together, this work establishes a biomaterial framework to support iCP differentiation and evaluates their performance relative to MSCs under defined 3D culture conditions for cartilage tissue engineering.

## 2. MATERIAL AND METHODS

### 2.1 Alginate gels fabrication and rheological analysis

Before use, unmodified (Sigma-Aldrich, UK, Cat. #W201502) and peptide-modified alginate powders were sterilized by exposure to 254 nm UV light a dose of 120,000 µJ/cm² for 30 minutes (CL-1000 Ultraviolet Crosslinker, Analytik Jena). Alginate solutions were prepared by dissolving sodium alginate powder in a 1:1 mixture of Dulbecco’s Phosphate-Buffered Saline without calcium and magnesium (DPBS^-/-^) and high-glucose DMEM (hgDMEM, Thermo-Fisher Scientific) to the desired final concentrations (1-3% w/v). Alginate hydrogel followed the protocol described by Sharma et al. [55], using a custom 5 cm × 10 cm mold designed to produce 6 discs, each 2 mm thick and 12.6 mm in diameter (**Figure S1A**).

Rheological assessments of cell-free alginate gels were conducted prior to immersion in cell culture medium (0h), and at subsequent time points under culture conditions. Briefly, following gelation, cell-free gels were maintained at 37°C and 5% CO₂ in hgDMEM with 1% penicillin/streptomycin (P/S) and 0.25 µg/mL Amphotericin B. Media was changed three times per week and gels were analyzed at 4 h, 24 h, 1 week and 2 weeks. Measurements were performed using an Anton Paar MCR-302 rheometer equipped with a 12.5 mm parallel plate geometry at room temperature (RT). During rheological analysis, sandpaper was used to prevent slipping of discs during measurements. Storage moduli (G′) and loss moduli (G″) were assessed within the linear viscoelastic region (LVR)⁶⁶. The amplitude sweep analysis was conducted with a constant normal force of 0.05 N [56]. Stress relaxation tests were performed on day 0 after an equilibration of gels in cell culture conditions for 24 hours. Gels underwent unconfined compression at a rate of 1 mm/min. Upon reaching 15% strain, the probe height was held constant while the normal force was recorded over time. The stress relaxation half time (τ½), an empirical measure of stress relaxation rate, was defined as the time required for the normal force to decrease to half of its initial value.

### 2.2 Peptide synthesis and alginate modification

Peptides (**Table 1**) were synthetized by solid-phase peptide synthesis (SPPS) on a microwave-assisted automatic synthesizer (LibertyBlue, CEM Corporations, USA), using a polystyrene (PS)-based resin (Rink Amide MBHA resin, Nova Biotech, USA) as solid support. Detailed protocol for peptide synthesis [57], purification by HPLC and analysis by LC-MS are provided in the **Supplementary Methods**.

**Table 1.** Peptide sequences and molecular weights. Molecular weights (g/mol) are reported as average (av.) and monoisotopic (mono.) values for each peptide.

| Peptide | Sequence | MW (g/mol) |
| --- | --- | --- |
| HAVDI | Ac-HAVDIGGGK-NH<sub>2</sub> | 893.891 (av.), 893.461 (mono.) |
| VAIDH | Ac-VAIDHGGGK-NH<sub>2</sub> | 893.891 (av.), 893.461 (mono.) |
| RGD | H-GGGGRGDSP-NH<sub>2</sub> | 757.77 (av.), 757.336 (mono.) |

For alginate modification, peptides were coupled to alginate chains using carbodiimide chemistry as previously described [58]. Briefly, 1-ethyl-3-(3-dimethylaminopropyl) carbodiimide hydrochloride (EDC; Fluorochem), *N*-hydroxysulfosuccinimide (sulfo-NHS; Fluorochem) and peptides (HAVDI/VAIDH: 120 mmol/g alginate, RGD: 13.3 mmol/g alginate) were added to a 1% (w/v) alginate solution in 0.1 M MES buffer and left to stir. After 24 hours at RT, the reaction was quenched with 10 mM hydroxylamine hydrochloride (Sigma Aldrich, UK). The solution was purified through extensive dialysis at room temperature using dialysis membranes (SpectraPor, UK, MWCO 12–14 kDa) against decreasing concentrations of NaCl in ultrapure deionized water until the conductivity was below 2 mS. Samples were then freeze-dried and stored at -20°C for up to 2 years. Peptide-modified alginate samples were then analyzed by nuclear magnetic resonance (NMR) and ^1^H diffusion-ordered spectroscopy (DOSY), as detailed in **Supplementary Methods**.

### 2.3 Human MSCs, iPSCs expansion and differentiation within gels

Bone marrow derived human MSCs (PromoCell GmbH, Cat. C-12974) were received at passage 1 and expanded in MSCs Expansion Medium (**Supplementary Methods**, **Table SM1**) at 37 °C with 5% CO_2_. Cells are routinely tested for mycoplasma. At passage 4, MSCs were seeded as single cells into alginate gels at a density of 5×10^6^ cells/mL. Following gelation, alginate discs were equilibrated in MSCs Expansion Medium for 24 hours before switching to chondrogenic media consisting of hgDMEM with GlutaMAX^TM^ (Thermo Fisher Scientific, UK) supplemented with 1% MEM non-essential amino acids (MEM-NEAA, Thermo Fisher, UK), 1% P/S, 0.25 μg/mL Amphotericin B, 1% ITS-Plus Premix (ITS+, Corning, NL), 6.6 nM dexamethasone (Sigma-Aldrich, UK), 100 μM ascorbate-2-phospate (A2P, Sigma-Aldrich, UK), 1 mM sodium-pyruvate, 40 μg/mL L-proline and 10 ng/mL human TGF-β3 (Peprotech, Thermo Fisher Scientific, UK). for 28 days. Gels maintained in MSCs Expansion Medium served as controls.

Human iPSCs (REPROCELL Europe Ltd, Cat. #802-3G) were received at passage 24 and were used at passage 28-30 for differentiation into chondroprogenitors. The cells received from REPROCELL were tested for mycoplasma, and their karyotype and pluripotency assessed prior to shipping. IPSCs were expanded on iMatrix-511-coated plates (Nippi, Inc., Cat. #NP892-011) with mTeSR Plus (MT+, STEMCELL Technologies, Cambridge, UK) at 37 °C with 5% CO_2_. iCP derivation was performed following the protocol of Dicks et al (2023) [33]. Briefly, iPSCs were differentiated to the chondroprogenitor stage through daily exposure to defined cocktails of growth factors and small molecules over 12 days. On day 12, cells were dissociated and analyzed through flow cytometry, used for chondrogenic differentiation, or stored in liquid nitrogen. ICP chondrogenic medium consisted of DMEM/F-12 (Thermo Fisher Scientific, UK) supplemented with 1% ITS+, 1% MEM-NEAA, 55 μM β-mercaptoethanol (Thermo Fisher Scientific, UK), 6.6 nM dexamethasone, 1% P/S, 50 μg/mL A2P, 40 μg/mL L-Proline, 10 ng/mL of human TGF-β3, 1 μM of Wnt-C59 (Axon Medchem, USA) and 1 μM of ML329 (Axon Medchem, USA). Chondrogenesis in 3D micromass culture was performed by centrifuging 0.5 × 10^6^ cells in iCP chondrogenic medium in 15 mL falcon tubes and kept for 28 days at 37 °C with 5% CO_2_. For chondrogenesis in alginate hydrogels, cells were resuspended as single cells in alginate solutions at a density of 20 × 10^6^ cells/mL, as at 5×10^6^ cells/mL no viable cells were observed. Crosslinked alginate discs were maintained for 28 days in iCP chondrogenic medium at 37 °C with 5% CO_2_. ICPs at day 12 of mesodermal differentiation served as a control due to the lack of expansion conditions capable of maintaining iCP phenotype [33].

### 2.4 Flow cytometry

Cells were prepared for flow cytometry by washing with FACS buffer, consisting of DPBS with 10% FBS, 0.1% (v/v) EDTA (0.5M stock, Thermo Fisher Scientific, UK) and 0.1% (w/v) sodium azide (Sigma-Aldrich, UK). Cells were then blocked with FcR Blocking Reagent (BioLegend, UK) and stained with specific antibodies (as listed in **Supplementary Methods**, **Table SM2**). After 30 min incubation, cells were washed, and fixed. Analysis was conducted using the BD LSR II cytometer with compensation performed using UltraComp eBeads (Thermo Fisher Scientific, UK). Viability staining was performed with Fixable Viability Dye eFluor 780 (Thermo Fisher Scientific, UK). Data was analyzed using FlowJo software (version 9.6.2, BD Biosciences, UK).

### 2.5 Viability assay

Cell viability within gels was assessed using the LIVE/DEAD Viability/Cytotoxicity Kit (Thermo Fisher Scientific, UK), according to manufacturer’s instructions. Gels were incubated with calcein-AM, ethidium homodimer and Hoechst dyes for 30 minutes at 37 °C with 5% CO_2_ before imaging with a Nikon Ti2 Eclipse confocal microscope. Viability was quantified from z-stack images and analyzed using Fiji software (version 2.14.0, imageJ), with values averaged across three independent regions per sample.

### 2.6 Gene expression analysis

For RNA extraction, hydrogel discs were washed with DPBS, homogenized in TRIzol, snap-frozen, and stored at -80°C. RNA was purified using a TRIzol/chloroform protocol, followed by isopropanol precipitation and further cleanup with RNeasy PowerClean Pro Kit (Qiagen, UK) according to the manufacturer’s instructions. Reverse transcription was performed using the High-Capacity cDNA Reverse Transcription Kit (Thermo Fisher Scientific, UK). Quantitative PCR (qPCR) was conducted on a QuantStudio 7 system using Power SYBR Green PCR Master Mix (Thermo Fisher Scientific, UK), according to manufacturer’s instructions. Primer sequences were obtained from Hodgkinson et al. (2021) [59] or designed in-house and purchased from Integrated DNA Technologies, as listed in **Supplementary Methods (Table SM3**).

### 2.7 Histological analysis

Hydrogels containing stem cells were fixed in 4% methanol-free formaldehyde (Thermo Fisher Scientific, UK) in HBSS with 7.5 mM CaCl_2_, processed, and embedded in paraffin. Sections were cut, mounted, and stained with alcian blue (Sigma Aldrich, UK) and fast nuclear red (Abcam, UK). For immunohistochemistry (IHC), sections underwent pepsin-based antigen retrieval (Thermo Fisher Scientific, UK), blocking, and staining with primary antibodies against collagen types I, II and aggrecan (**Supplementary Methods**, **Table SM4**), followed by secondary antibody staining and 3,3′-diaminobenzidine (DAB) chromogen application. Slides were counterstained with hematoxylin, dehydrated, and mounted with DPX (Sigma Aldrich, UK) before imaging.

### 2.8 Statistical analysis

Details of statistical analyses, including the specific tests used and sample sizes, are provided in individual figure legends. Statistical analyses were performed using GraphPad Prism 9 software (GraphPad). Statistical significance was determined at a threshold of P < 0.05 (**Supplementary Methods**, **Table SM5**).

## 3. RESULTS

### 3.1 Ionically Crosslinked 1–3% Unmodified Alginate Gels Exhibit Increasing Stiffness and Fast Stress Relaxation

Although the influence of biomaterial stiffness on the fate of stem cells being well-established, the optimal stiffness for chondrogenesis may depend on both the biomaterial composition and the stem cells used [45,46,60–63]. Alginate gels were generated at varying polymer concentrations (1%, 2%, and 3% w/v) to explore a range of stiffnesses previously associated with chondrogenic differentiation [45,60–64]. The resulting gels, shaped as 2 mm-high, 12 mm-diameter discs, exhibited increasing opacity with higher concentrations (**Figure 1A**). Consistent with previous reports [65], the storage modulus (G′), indicative of gel stiffness, increased proportionally with alginate concentration: after 24 hours in culture medium, 1%, 2%, and 3% (w/v) gels showed G′ values of 0.37 ± 0.05 kPa, 2.17 ± 0.32 kPa, and 4.55 ± 0.52 kPa, respectively (**Table 2**).

**Figure 1.**
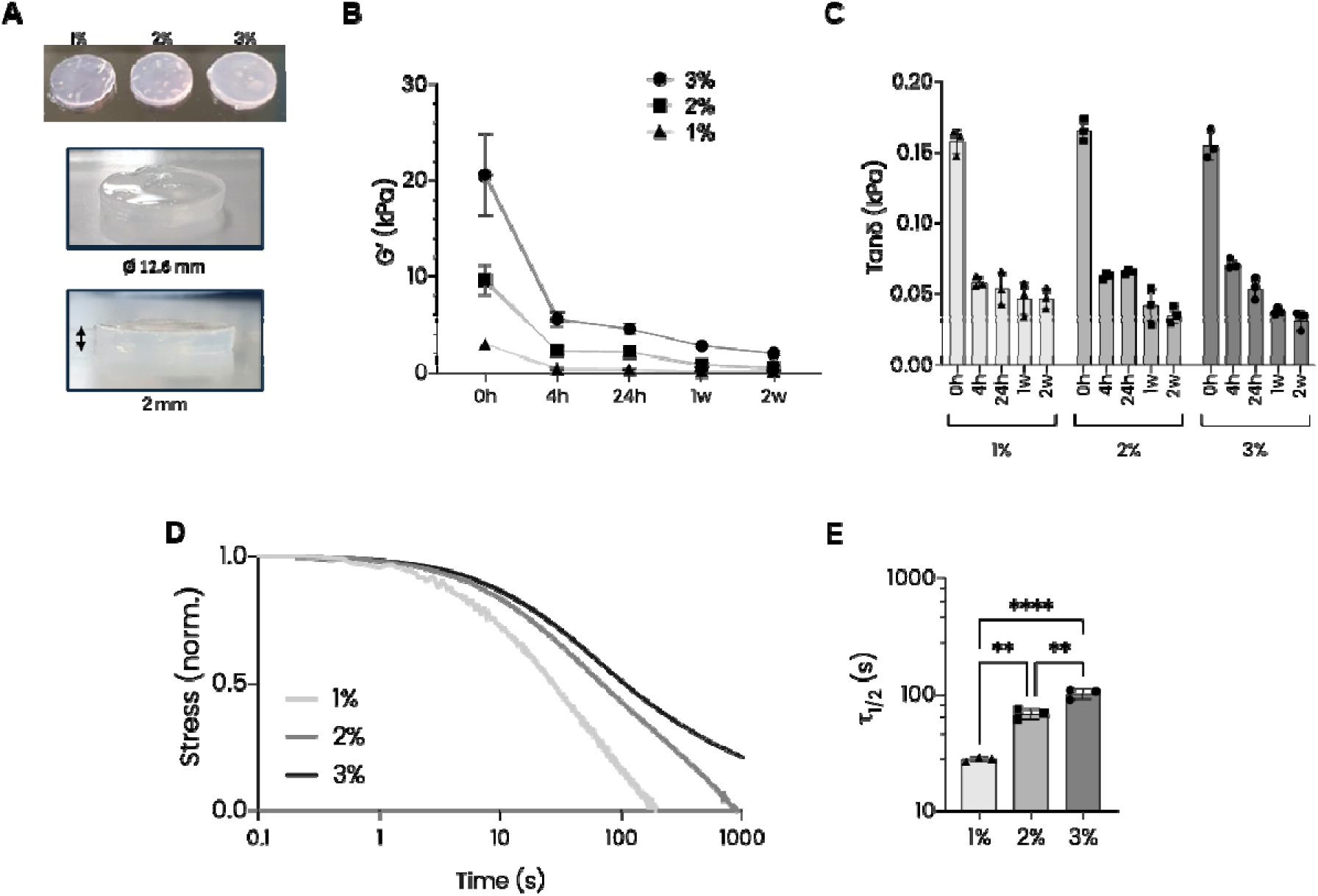
Rheological analysis of 1-3 % (w/v) alginate hydrogels in cell culture medium. **A**) Representative images of 1-3% (w/v) gels. Hydrogels dimensions were approximately 2 mm in thickness, and 12.6 mm in diameter. **B**) Stiffness measured as storage modulus (G’) analyzed through amplitude sweep, with values stabilizing after 1-2 weeks under cell culture conditions. **C**) Tanδ measurement indicated that alginate gels became more elastic over time, with limited variation across different stiffness conditions. Statistical analysis was performed using two-way ANOVA with Tukey’s multiple comparisons for normally distributed data, and Mann-Whitney with Holm-Šídák correction for not-normally distributed data. **D**) Stress relaxation test on alginate hydrogels after 24 hours in growth medium (15% strain; normalized data). **E**) Quantification of relaxation timescale from (D), showing the time required for gels to relax to half of their initial stress (*τ*_1/2_). One-way ANOVA with Tukey’s multiple comparisons was performed. Error bars and symbols represent mean ± s.d. (n = 3).

**Table 2.** G’ values of 1-3% (w/v) alginate gels measured after 24 h in cell culture medium. These values are used hereafter as reference stiffness for each concentration (mean ± s.d, n=3).

| Alginate concentration | 1% (w/v) | 2% (w/v) | 3% (w/v) |
| --- | --- | --- | --- |
| Stiffness at 24 h (kPa) | $0.37 \pm 0.05$ | $2.17 \pm 0.32$ | $4.55 \pm 0.52$ |

All gels softened over time, likely due to ion exchange with the culture media [66]. However, values stabilized after 1 week of culture (**Figure 1B**). Alginate discs maintained a gel-like behavior (G′> G″) at all time points (**Figure S1B**). Tanδ values initially displayed significant differences among gel concentrations, but they became comparable between percentages after 24 hours of equilibration in medium (**Figure 1C**). Over time, all gels exhibited a progressive decline in tanδ, indicating a shift toward increased elasticity and reduced viscous dissipation. This trend stabilized after one week of incubation, mirroring the pattern observed for storage modulus. Although an elastic-dominant profile is characteristic of ionically crosslinked alginates, highly elastic environments have been linked to reduced N-cadherin expression and impaired chondrogenic potential in MSCs [48].

Given the established role of stress relaxation in regulating stem cell behavior [48,63,67], rheological analysis was performed to evaluate stress relaxation across all alginate gel concentrations. As shown in **Figure 1D-E**, 1% and 2% (w/v) alginate gels demonstrated rapid stress relaxation, with τ₁/₂ values of less than one minute. Moreover, these gels reached complete relaxation in less than 15 minutes, a timeframe typical of fast relaxing gels previously shown to promote chondrogenesis [63,67]. Conversely, the 3% gel exhibited a slower relaxation profile, with a τ₁/₂ exceeding one minute, and required more time to completely relax under stress.

Overall, these findings established alginate hydrogels spanning 0.37–4.55 kPa with physiologically relevant stress relaxation properties suitable for chondrogenic differentiation studies. Having established this mechanically tunable platform, we next sought to determine how substrate stiffness influences the chondrogenic differentiation of MSCs, providing a baseline for subsequent comparison with iPSC-derived chondroprogenitors.

### 3.2 Unmodified Alginate Stiffness Influences Chondrogenic Differentiation of MSCs

To assess the impact of alginate hydrogel mechanics on chondrogenesis, human MSCs were embedded as single cells in unmodified 1-3% (w/v) alginate gels and cultured in either chondrogenic or expansion medium for 28 days. At early time points, MSCs in chondrogenic medium showed high viability across all gel concentrations, with approximately 90% viable cells by day 7 (**Figure 2A**). There was no significant difference in viability among the gels, indicating that differences in stiffness did not affect early cell viability. Morphologically, MSCs retained a rounded phenotype across all gel conditions during the first seven days of culture (**Figure 2A**).

**Figure 2.**
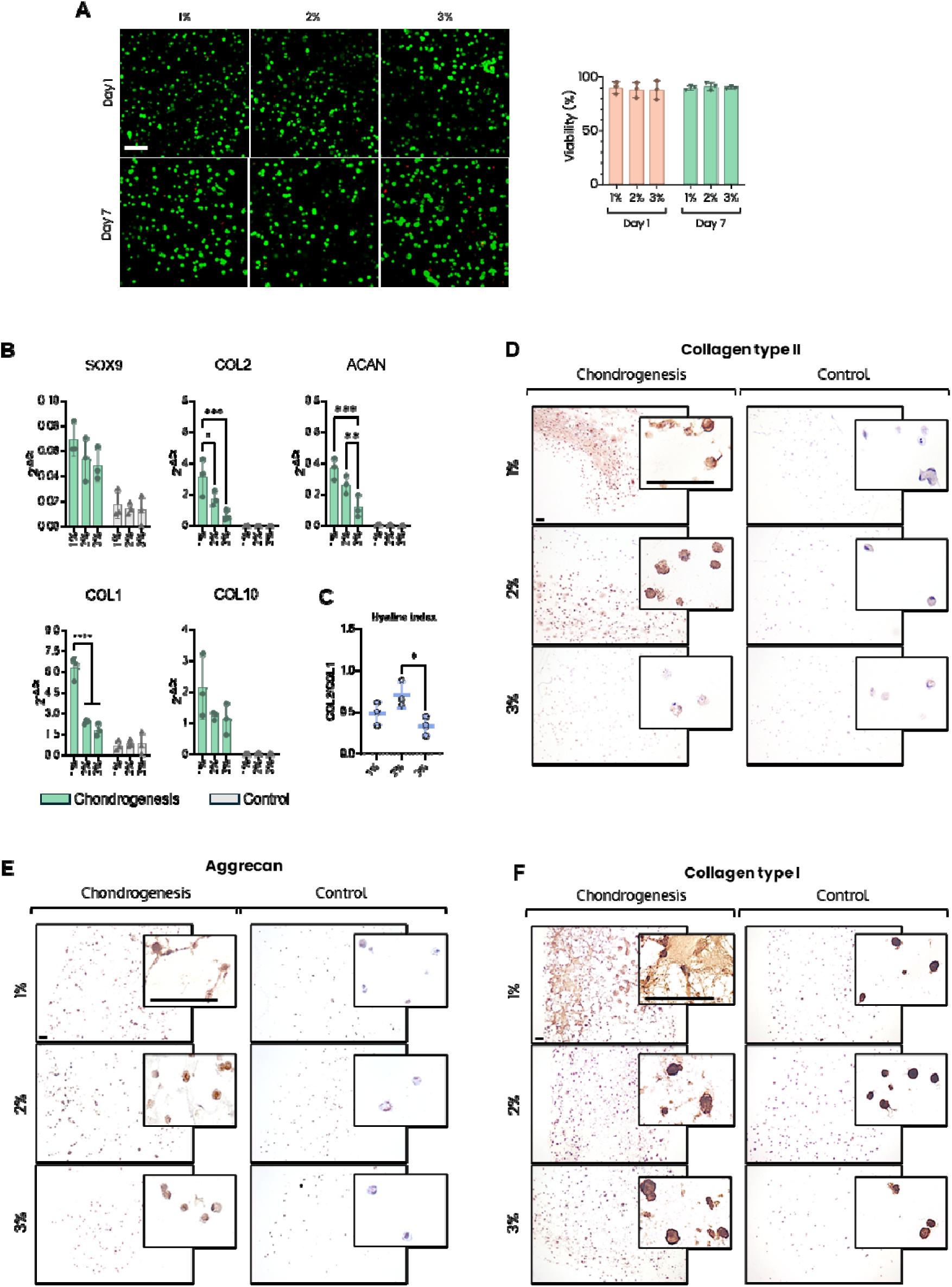
Chondrogenesis of MSCs in 1-3% (w/v) alginate gels. **A**) Viability of MSCs in 1-3% (w/v) gels assessed by Live/Dead assay. Representative images (left) and quantification (right) at day 1 and day 7 of chondrogenesis are shown. Two-way ANOVA with Tukey’s multiple comparisons post hoc test was performed. Error bars represent mean ± s.d. (n = 3). Green: viable cells (calcein-AM), red: dead cells (ethidium homodimer-1). Scale bar = 100 μm. **B**) Expression of chondrogenic markers of MSCs embedded in 1-3% (w/v) alginate gels after 28 days in chondrogenic or control medium. Statistical analysis was performed using two-way ANOVA with Tukey’s multiple comparisons (Kruskal-Wallis when data not normally distributed) post hoc test was performed. Error bars and symbols represent mean ± s.d. (n = 3). **C**) COL2/COL1 ratio of MSCs on day 28 of chondrogenesis. One-way ANOVA with Tukey’s multiple comparison post hoc test was performed. Error bars and symbols represent mean ± s.d. (n = 3). **D-F**) Immunohistological staining of ECM proteins in hMSCs cultured in 1-3% (w/v) alginate gels after 28 days of chondrogenic differentiation. Paraffin-embedded sections were stained for (**D**) collagen type II, (**E**) aggrecan and (**F**) collagen type I. Antibody specificity was validated using isotype control and positive controls (**Figure S4**). Experiments were repeated three times with comparable results. Scale bar = 100 μm.

Gene expression analysis after 28 days revealed distinct differences in chondrogenic marker expression across the gel concentrations (**Figure 2B**). MSCs cultured in 1% and 2% gels exhibited significantly increased expression of collagen type II (COL2) and aggrecan (ACAN), key components of hyaline cartilage, compared to those in 3% gels. In contrast, SOX9 expression, a transcription factor critical for chondrogenesis, was comparable across percentages. Notably, expression of collagen type I (COL1), commonly associated with fibrocartilage, was markedly higher in the 1% (w/v) gel than in the 2% and 3% (w/v) gels. This observation was reflected in the hyaline index (COL2/COL1 ratio), which was highest in the 2% gels (**Figure 2C**), indicating a more hyaline-like cartilage phenotype. Collagen type X (COL10), a marker of hypertrophy, was detected across all conditions at comparable levels, suggesting that MSCs expressed this hypertrophy marker regardless of gel stiffness. Immunohistochemical (IHC) analysis (**Figure 2D-E**) confirmed deposition of collagen type II and aggrecan most prominently in the 1% and 2% gels. Glycosaminoglycan deposition was further confirmed through alcian blue staining (**Figure S2A**). In the 1% gels, collagen type I deposition (**Figure 2F)** was notably elevated and accompanied by the presence of elongated cells, whereas a rounded morphology was maintained in all other conditions (**Figure S2B-C**). These morphological characteristics resemble features previously reported in dedifferentiated chondrocytes [68–70] and in degenerative cartilage regions [71,72], potentially explaining the increased collagen type I production observed in the softest gels.

Together, these results identify 2% alginate as a mechanically favorable condition for chondrogenesis of MSCs, while also highlighting persistent limitations in achieving a stable hyaline cartilage phenotype, thereby motivating evaluation of alternative cell sources.

### 3.3 Unmodified Alginate Gel Stiffness Influences Chondrogenic Differentiation of iPSC-Derived Chondroprogenitors

Given the limitations observed in hMSCs-derived cartilage, we next evaluated whether iPSC-derived chondroprogenitors (iCPs) could survive and differentiate within alginate hydrogels, and whether they would exhibit enhanced chondrogenic potential within a specific biomaterial stiffness. iCPs were generated following a previously established differentiation protocol [33] (**Figure 3A**): immunophenotypic validation at day 12 showed that approximately 25% of cells displayed a CD146^+^CD166^+^CD44^+^CD34^−^CD45^−^ phenotype (**Figure 3B**), consistent with prior reports [33,39,40,73,74]. Notably, over 50% of cells expressed CD166, a marker delineating progenitor cells with strong chondrogenic potential [40,75]. While this represents an heterogeneous population, chondrogenic differentiation was performed following the established protocol [33], which includes medium supplementation with Wnt-C59 and ML329 inhibitors to suppress off-target differentiation and enrich for the CP subpopulation. To confirm their identity and demonstrate their chondrogenic ability, iCPs were differentiated as pellet cultures for 28 days (**Figure 3C**). At the end of chondrogenesis, pellets composition was comparable with what reported by the authors of the protocol: immunohistochemical staining (**Figure 3D**) revealed collagen type II and aggrecan within the pellets’ core, while collagen type I was limited to the outer layers, resembling a perichondrium-like structure. The substantial glycosaminoglycan deposition was further confirmed through strong alcian blue staining (**Figure S2D**).

**Figure 3.**
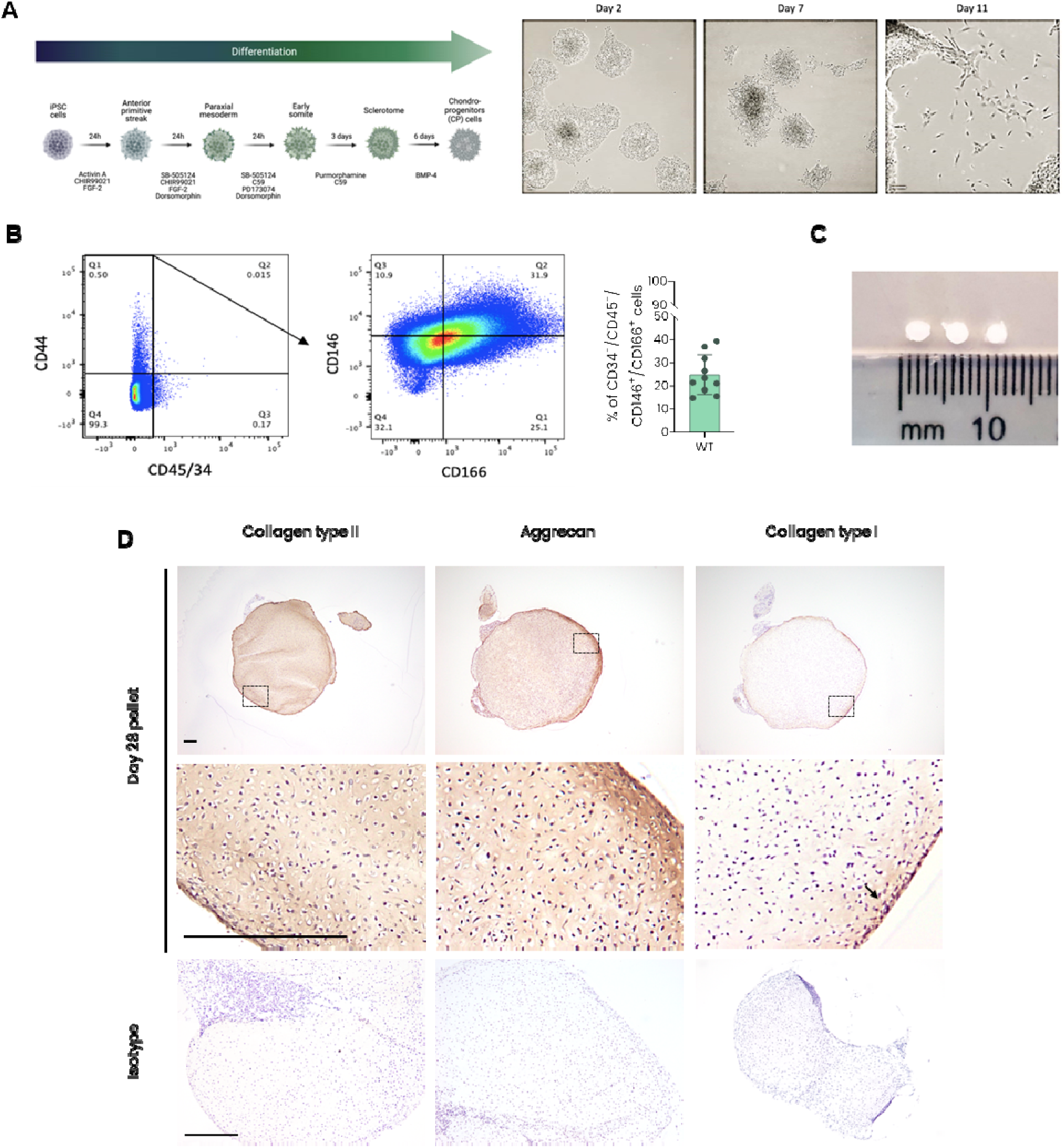
Derivation of iCPs and chondrogenic pellet analysis. **A**) Schematic representation of stepwise mesodermal differentiation of human iPSCs towards chondroprogenitor-like cells (right), with corresponding bright-field images showing cell morphology at different time points (left). During differentiation, elongated cells emerge from iPSCs colonies and reach a spindle-shaped morphology by day 12. This morphology is consistent with previously reported differentiation protocols (i.e., Dicks et al., *Methods Mol Biol*, 2598, 2023). Created with BioRender.com. **B**) Flow cytometry gating strategy used to quantify chondroprogenitor populations (left) and corresponding percentage of CD34-/CD45-/CD146+/CD166+ iCPs at day 12 (right). Error bars represent mean ± s.d. (n = 10). **C**) Representative images of pellets after 28 days of chondrogenic differentiation. **D**) Representative IHC of iCPs pellets after 28 days, showing expression of collagen type II, aggrecan, collagen type I, with hematoxylin counterstaining to visualize nuclei. Scale bars = 100 μm.

Having confirmed their chondrogenic potential, iCPs at day 12 of mesodermal differentiation were lifted, embedded as single cells in 1-3% alginate gels and cultured for 28 days in chondrogenic medium. At day 1 and 7 of differentiation, gels were analyzed to verify early viability and overall morphology. Embedded iCPs showed reduced viability by day 7, with approximately 16 to 18% viable cells remaining (**Figure 4A**). Although not directly investigated, this low viability may reflect selection of a specific chondrogenic subpopulation, potentially driven by the off-target inhibitors present in the differentiation medium, similar to observations previously reported in pellet cultures [39], and by the biomaterial’s properties. Interestingly, in contrast to hMSCs, iCPs exhibited clustering behavior, with viable cells often observed within multicellular aggregates at day 7. However, isolated cells frequently appeared nonviable, suggesting that cluster formation itself may further contribute to the selective survival of a specific iCP subpopulation within the gels.

**Figure 4.**
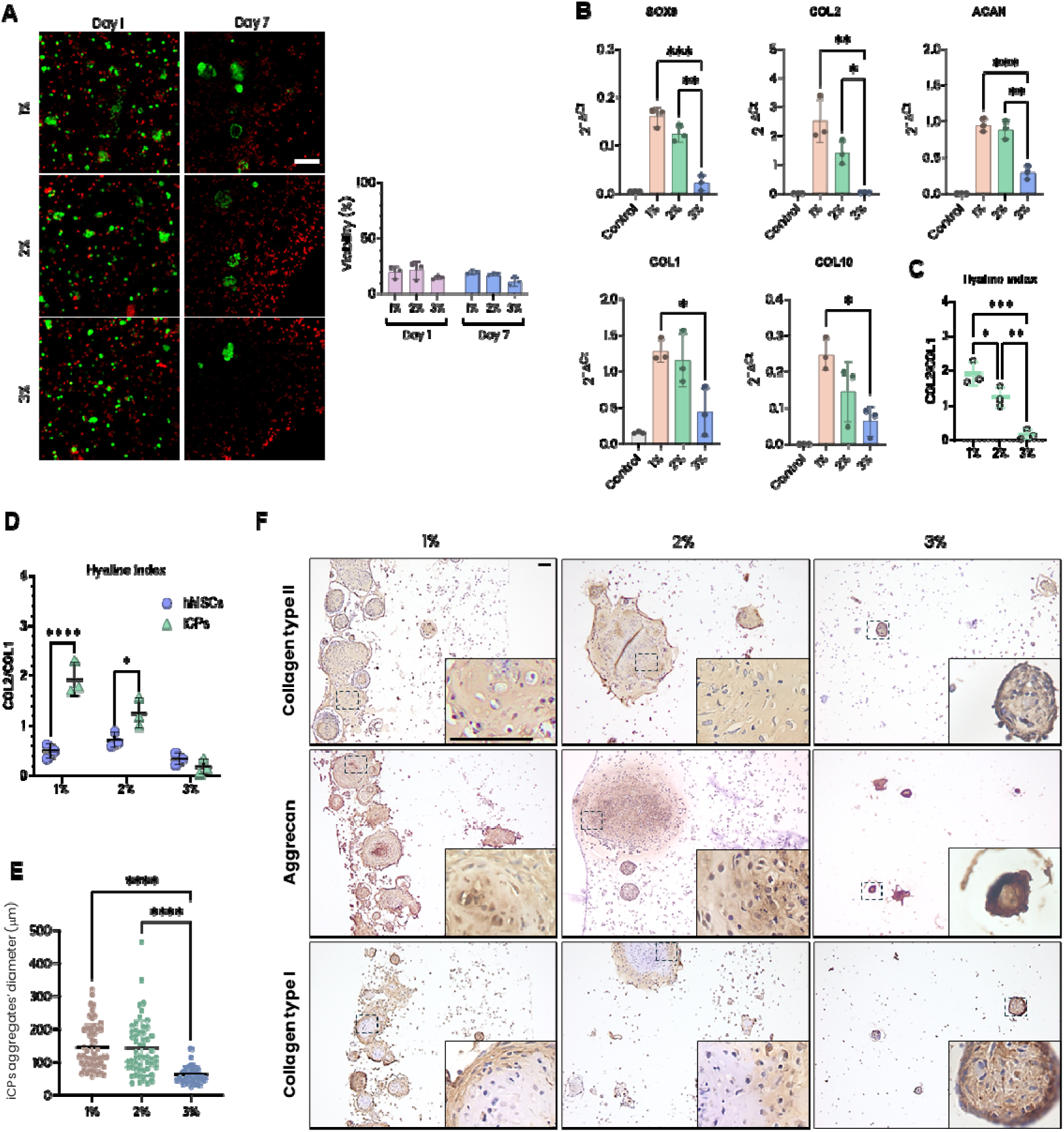
Chondrogenesis of iCPs in 1-3% (w/v) gels. **A**) Viability of iCPs in 1-3% (w/v) gels assessed by Live/Dead assay. Representative images (left) and quantification (right) at day 1 and day 7 of chondrogenesis are shown. Two-way ANOVA with Tukey’s multiple comparisons post-hoc test was performed. Error bars represent mean ± s.d. (n = 3). Green: viable cells (calcein-AM), red: dead cells (ethidium homodimer-1). **B**) Expression of chondrogenic markers in iCPs embedded in 1-3% (w/v) alginate gels after 28 days of differentiation, determined by qPCR. One-way ANOVA with Tukey’s multiple comparisons post-hoc test was performed. Error bars represent mean ± s.d. (n = 3). **C**) COL2/COL1 ratio of iCPs on day 28 of chondrogenesis. One-way ANOVA with Tukey’s multiple comparisons post-hoc test was performed. Error bars represent mean ± s.d. (n = 3). **D**) Comparison of COL2/COL1 ratios between hMSCs and iCPs on day 28 of chondrogenesis. Statistical analysis was performed using an unpaired t-test (n=3). **E**) Quantification of pellet diameter measured from histological sections. Statistical analysis was performed using one-way ANOVA or Kruskal-Wallis test, as appropriate (n=49-54 from two sections per sample across the three biological replicates). Error bars represent mean ± s.d. (n = 3). **F**) Immunohistological staining for ECM proteins in iCPs cultured in 1-3% (w/v) alginate gels for 28 days under chondrogenic conditions. Paraffin-embedded sections were stained for collagen type II, aggrecan, and collagen type I. Experiments were repeated three times with comparable results. Scale bars = 100 μm.

Gene expression profiling at the end of differentiation (day 28) indicated that softer gels (1% and 2%) significantly enhanced chondrogenic marker expression compared to the 3% gels. Specifically, expression of SOX9, COL2, ACAN and COL1 was significantly elevated in the 1% and 2% gels compared to 3% gels (p < 0.05; **Figure 4B**). In contrast to hMSCs, COL10 levels were also elevated in 1-2% gels relative to the 3% gel. However, the hyaline index was highest in the 1%, with 2% gels right below, and 3% gels showing the lowest index, further supporting the notion that the softer matrices fostered a hyaline cartilage phenotype more effectively than the 3% gel (**Figure 4C**). Notably, iCPs in softer gels exhibited a higher hyaline index than hMSCs (**Figure 4D**), along with increased ACAN expression, moderately decreased COL1 expression, and markedly lower COL10 expression than hMSCs (**Figure S2E**). Histological analysis revealed that iCPs did not differentiate as single cells, as previously observed with MSCs, but rather formed well-defined chondrocytic aggregates within 1% and 2% gels. Interestingly, these aggregates were markedly smaller in the stiffer 3% gels (**Figure 4E**). Immunohistochemical analysis confirmed collagen type II and aggrecan deposition across all conditions (**Figure 4F**), and proteoglycan deposition was further confirmed through alcian blue staining (**Figure S2F**). Interestingly, in 1-2% gels collagen type I was observed primarily localised in the outer layers of the aggregates, whereas in 3% gels it appeared more uniformly distributed (**Figure 4F**).

These findings reveal divergent responses between hMSCs and iCPs in unmodified alginate: whilst hMSCs differentiated as single cells, iCPs preferentially formed cartilaginous aggregates in softer gels and showed higher hyaline indices and proteoglycan deposition compared to hMSCs. Critically, 2% gels proved optimal for both cell types: in hMSCs, they yielded the highest hyaline index whilst avoiding collagen type I overexpression observed in 1% gels; in iCPs, they supported robust aggregate formation with high hyaline index. Thus, this stiffness was selected for peptide functionalization studies.

### 3.4 Peptide Functionalization Differentially Affects Viability, Chondrogenic Gene Expression, and ECM Composition in MSCs and iCPs

Having identified 2% gels as having optimal stiffness, we next investigated whether incorporating bioactive peptides could further enhance chondrogenesis by modulating cell-matrix and cell-cell interactions. To better replicate the microenvironment of cartilage development and enhance chondrogenesis, alginate gels were functionalized with RGD (arginine-glycine-aspartic acid) and HAVDI (histidine-alanine-valine-aspartic acid-valine) peptides to mimic extracellular matrix adhesion and N-cadherin-mediated interactions, respectively. Incorporation of these peptides has proved beneficial for chondrogenic differentiation of human MSCs [76–79].The RGD peptide, derived from fibronectin, is widely used to mimic cell-ECM interactions and promote adhesion. Moreover, fibronectin is a critical ECM component during mesenchymal condensation and growth plate development[80,81]. However, previous studies have shown variable effects of RGD on stem cell chondrogenesis [82–88], potentially reflecting differences in concentration, timing, or culture conditions. HAVDI, originating from the extracellular domain of N-cadherins, serves to simulate cell-cell interactions. Thus, peptides GGGGRGDSP, HAVDIGGGK, and a scrambled form of HAVDI (VAIDHGGGK) were synthesized and covalently attached to alginate. Peptides were initially synthesized, and purity was confirmed through analytical HPLC (**Figure S3A**). Covalent bonding between peptides and alginate chains was confirmed through both ^1^H NMR and DOSY ^1^H NMR (**Figure S3B-D**). Peptide incorporation did not significantly alter hydrogels’ mechanics, with only a modest increase in viscosity observed in HAVDI/RGD gels after prolonged culture (**Figure S4**).

To assess how peptide functionalization differentially influences MSCs and iCPs, both cell types were encapsulated in 2% peptide-functionalized alginate gels.

At early timepoints MSCs exhibited excellent viability (approximately 90%), with no significant decline observed by day 7. However, RGD inclusion induced some cells to adopt a fibroblastic shape by day 7, whereas cells in unmodified gels retained their round phenotype. Despite these morphological differences, overall MSC viability remained comparable between functionalized and unmodified alginate gels (**Figure 5A**), indicating that peptide incorporation does not compromise cell survival but may influence cell morphology at early time points.

**Figure 5.**
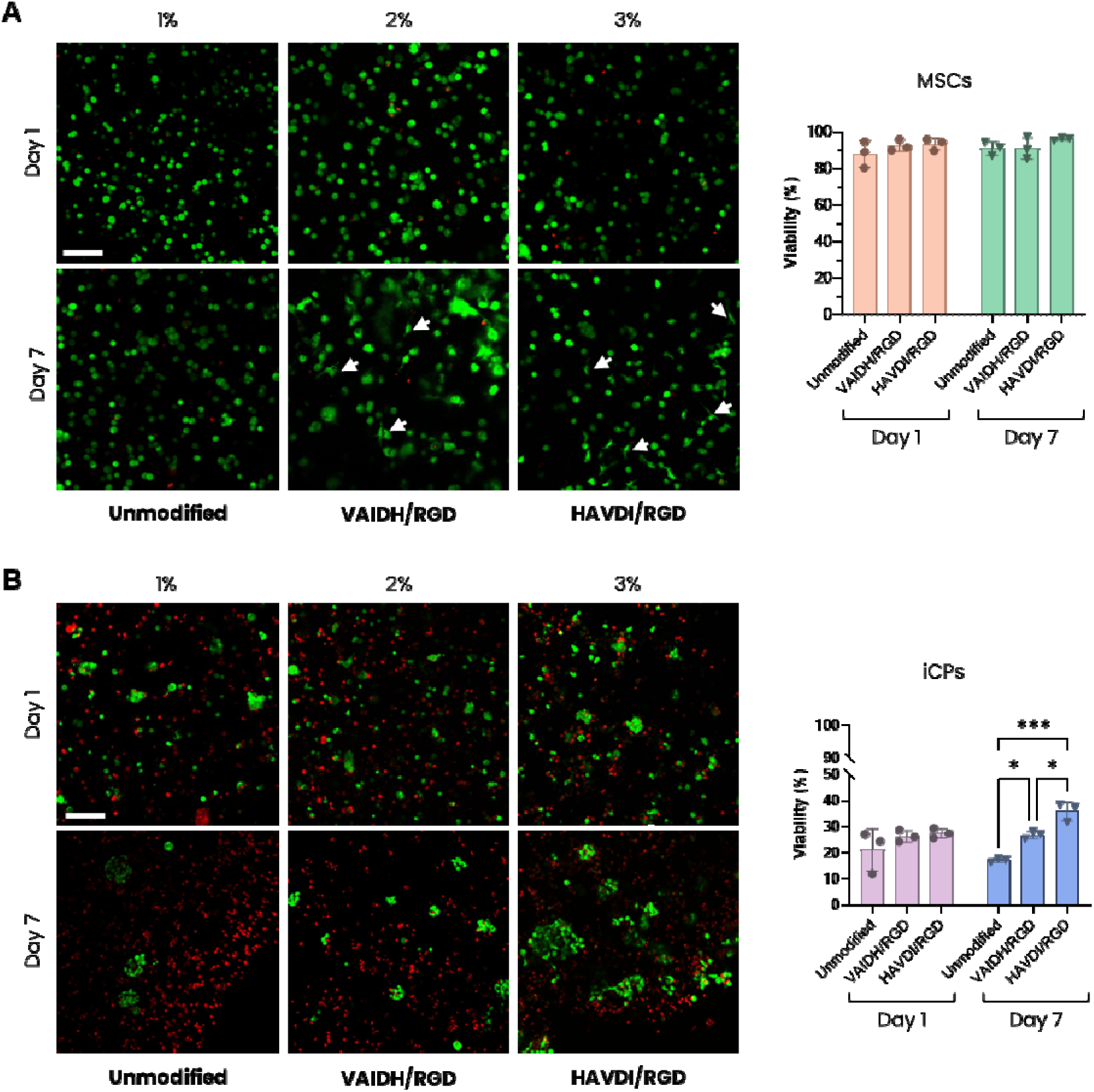
Viability of MSCs and iCPs in peptide-functionalized gels. Representative images and quantitative analysis of cell viability assessed by Live/Dead Assay for MSCs (**A**) and iCPs (**B**) at day 1 and day 7 of chondrogenesis. White arrows indicate cells showing elongate morphologies. Two-way ANOVA with Tukey’s multiple comparisons post hoc test was performed. Error bars represent mean ± s.d. (n = 3). Green: viable cells (calcein-AM), red: dead cells (ethidium homodimer-1). Scale bar = 100 μm.

For iCPs, initial viability was similar in both non-modified and peptide-functionalized alginate gels (**Figure 5B**), indicating that the presence of peptides did not significantly affect cell viability within the first 24 hours after seeding. By day 7, however, viability increased in both VAIDH/RGD and HAVDI/RGD functionalized gels compared to non-modified controls. Cells in VAIDH/RGD gels formed compact aggregates similar to those observed in unmodified gels, though these aggregates appeared more abundant in the presence of RGD. In contrast, cells in HAVDI/RGD gels formed larger but less compact clusters. Whether these larger structures resulted from cell proliferation or migration of viable cells was not further investigated. Moreover, unlike MSCs, iCPs consistently retained their rounded morphology for the first 7 days, with no evidence of elongation (**Figure 5B**), indicating a different response to integrin-binding ligands.

To determine how these early differences in cell organization impact chondrogenesis, both hMSCs and iCPs were cultured in peptide-functionalized alginate gels and analyzed at day 28 of differentiation. qPCR analysis of hMSCs demonstrated that incorporation of RGD into the gels significantly upregulated COL1 expression while markedly reducing COL10 levels (**Figure 6A**). In contrast, COL2 and ACAN expression showed modest but non-significant increases. Notably, the addition of HAVDI further enhanced COL2 and ACAN expression but had no significant effect on COL10 or COL1 levels compared to RGD-only gels, while SOX9 expression remained consistent across all conditions (**Figure 6A**). Consistent with these findings, the absence of HAVDI was associated with a lower hyaline index (**Figure 6B**). Histological analysis corroborated the qPCR findings, demonstrating substantially increased deposition of collagen type II and I, along with moderately enhanced aggrecan deposition in HAVDI/RGD gels compared to unmodified gels (**Figure 6C-E**).

**Figure 6.**
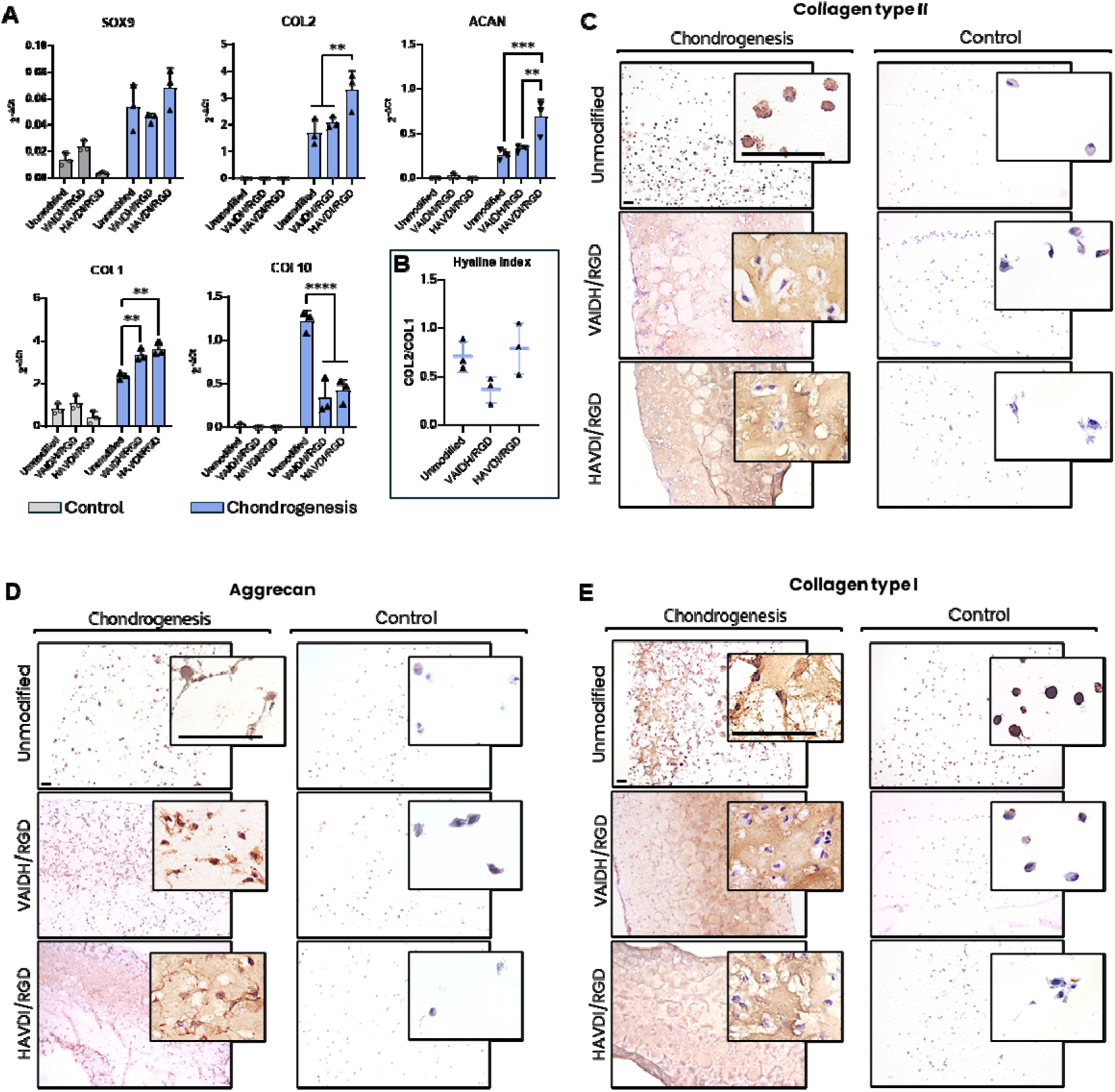
Chondrogenesis of MSCs in peptide-functionalized alginate gels. **A**) Expression of chondrogenic markers in hMSCs embedded in unmodified and peptide-functionalized gels after 28 days in chondrogenic or control medium. Statistical analysis was performed using one-way ANOVA with Tukey’s multiple comparison post-hoc test was performed. Error bars represent mean ± s.d. (n = 3); **B**) Hyaline Index of hMSCs in unmodified and peptide-functionalized gels after 28 days of chondrogenesis. Error bars represent mean ± s.d. (n = 3). **C-E**) Immunohistological staining for ECM proteins in hMSCs cultured in unmodified and peptide-functionalized gels after 28 days of chondrogenic differentiation. Paraffin-embedded sections were stained for (**C**) collagen type II, (**D**) aggrecan, and (**E**) collagen type I. Experiments were repeated three times with comparable results. Scale bar = 100 μm.

In contrast, transcript analysis of iCPs at day 28 showed that incorporation of RGD moderately reduced hyaline cartilage markers (COL2, ACAN) compared to unmodified gels, while significantly increasing COL1 and suppressing COL10 expression (**Figure 7A**). The addition of HAVDI to RGD-functionalized gels significantly enhanced SOX9, COL2 and ACAN expression, but did not have a significant effect on COL1 or COL10 levels compared to RGD-only gels. Moreover, VAIDH/RGD gels showed a significantly lower hyaline index than HAVDI/RGD gels (**Figure 7B**). Interestingly, when comparing gene expression between the two cell types, iCPs in HAVDI/RGD gels exhibited reduced COL10 expression and increased levels of ACAN and SOX9 compared to MSCs (**Figure 7C-D**) but showed comparable hyaline index. In the absence of HAVDI, iCPs still displayed significantly lower COL10 expression relative to MSCs but showed no notable differences in hyaline cartilage gene expression. To further assess hypertrophic potential, MMP13, RUNX2 and ALP expression were additionally examined alongside COL10 in both VAIDH/RGD and HAVDI/RGD gels (**Figure 7C**). Consistent with the COL10 findings, MMP13 expression was significantly lower in iCPs compared to hMSCs in both conditions. In contrast, RUNX2 and ALP expression did not differ significantly between cell types in either condition.

**Figure 7.**
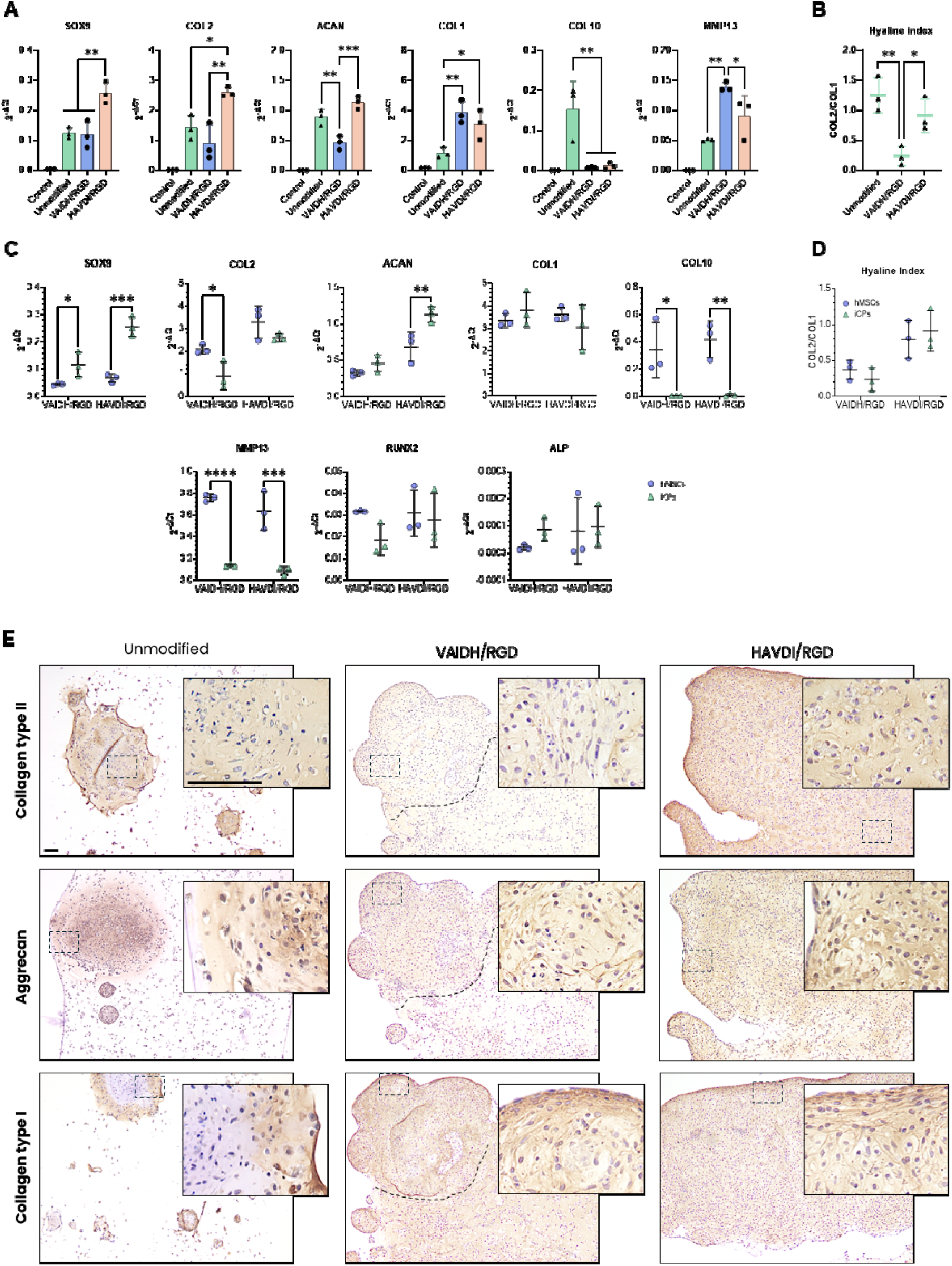
Chondrogenesis of ICPs in peptide-functionalized alginate gels. **A**) Expression of chondrogenic markers in iCPs embedded in unmodified and peptide-functionalized gels after 28 days of differentiation, determined by qPCR. One-way ANOVA with Tukey’s multiple comparison post-hoc test was performed (n=3). Error bars represent mean ± s.d. (n = 3). **B**) Hyaline index of cells cultured in unmodified and peptide-functionalized gels after 28 days of chondrogenesis. One-way ANOVA with Tukey’s multiple comparison post-hoc test was performed (n=3). **C**) Gene expression and **D**) hyaline index comparison between hMSCs and iCPs cultured in peptide-functionalized gels after 28 days of chondrogenesis. Statistical analysis was performed using multiple t-test with Holm-Šídák correction for multiple comparisons. Error bars represent mean ± s.d. (n = 3). **E**) Immunohistological staining for ECM proteins in iCPs cultured in unmodified and peptide-functionalized gels were stained for each Paraffin-embedded sections were stained for collagen type II, aggrecan, and collagen type I. The dashed line indicates regions of higher cell density. Experiments were repeated three times with consistent results. Scale bar = 100 μm.

Histological analysis showed that functionalized gels promoted increased ECM deposition and a more uniform cell distribution compared to non-functionalized gels (**Figure 7E**). In VAIDH/RGD gels, structures of higher cell density, were observed (highlighted by dashed lines, **Figure 7E**). Whether these result from the fusion or the expansion of smaller aggregates identified during viability assessments was not investigated. Within these gels, collagen type II was evenly distributed within the body of the section, while aggrecan was predominantly localized to denser aggregates and less expressed in less populated areas. HAVDI integration visibly enhanced collagen type II and aggrecan protein expression, consistent with transcript data. Interestingly, in presence of HAVDI both cell distribution and ECM proteins deposition were evenly distributed throughout the gel (**Figure 7E**). Collagen type I also remained uniformly present in the body of the gel sections. Notably, cells at the periphery of the gels in both functionalized conditions displayed a flattened morphology, resembling the superficial zone of native articular cartilage[89].

Together, these findings demonstrate that peptide functionalization differentially modulates MSC and iCP chondrogenesis, with RGD alone promoting fibrocartilage markers in both cell types, while the HAVDI/RGD combination significantly enhances hyaline cartilage matrix deposition. Notably, iCPs maintained a more favorable chondrogenic profile than hMSCs, exhibiting lower COL10 expression and higher SOX9 and ACAN levels in HAVDI/RGD-functionalized gels.

## 4. DISCUSSION

MSCs have been widely used for cartilage modelling and tissue engineering; however, their tendency to form fibrocartilage, undergo hypertrophic differentiation, and exhibit limited proliferative capacities constrains their broader utility [13–15,22]. Chondroprogenitor cells have emerged as a promising alternative, as they differentiate into chondrocytes with reduced hypertrophy and the greater capacity to generate hyaline cartilage [26,27]. In this study, we directly compared these two cell types within defined biomaterial environments to determine how cell-intrinsic properties and matrix cues interact to regulate chondrogenesis, with the aim of improving biomaterial-guided cartilage modelling in 3D culture systems.

First, we fabricated hydrogel discs with stiffness ranging from 0.37 to 4.55 kPa (**Table 1**) and physiologically relevant stress relaxation properties to support chondrogenic differentiation of stem cells and ECM deposition [45,60–64,90]. Chondrogenic differentiation within these non-functionalized gels revealed marked differences in cell behavior between iCPs and hMSCs. For hMSCs, the softest (1%) gels promoted an elevated collagen type I expression and the emergence of elongated cell morphologies. This elongated morphology may reflect reduced mechanical confinement in softer gels, potentially facilitating cytoskeletal tension and stress fiber formation [66,91]. Such changes have previously been associated with dedifferentiation or fibrocartilage-like phenotypes [68,69,71,92–94]. The stiffest (3%) gels suppressed hyaline cartilage markers (collagen type II and aggrecan), but intermediate stiffness (2%) provided an optimal mechanical environment, supporting robust expression of hyaline cartilage markers while controlling the expression of collagen type I. Interestingly, while collagen type II, type I and aggrecan expression was significantly affected by stiffness, COL10A1 expression remained relatively stable across all conditions. Both fibrocartilage tendency (COL1) and hypertrophic differentiation (COL10) are commonly detected during hMSCs chondrogenesis [13,21,22,42,95], suggesting that fibrocartilage-associated and hypertrophic differentiation programmes in hMSCs might be differentially regulated by mechanical cues.

iCPs exhibited distinct behavior from hMSCs in unmodified alginate gels. While hMSCs survived and deposited matrix as single cells, iCPs showed limited initial survival and instead formed cell clusters, despite being seeded as single cells. This clustering likely reflects either aggregation of neighboring cells or proliferation of surviving cells, and resembles mesenchymal condensation during embryonic chondrogenesis, where close cell-cell contact is required for lineage commitment and cartilage formation [96–100]. The reduced early viability of iCPs relative to hMSCs may reflect selective enrichment for chondrogenically competent cells: the day-12 iCP population is inherently heterogeneous, retaining off-target cells alongside those with chondrogenic potential, and the combination of off-target inhibitors in the differentiation medium and the biomaterial environment likely eliminated non-chondrogenic cells while surviving cells successfully formed cartilaginous aggregates. Single-cell encapsulation was selected as the seeding strategy for both cell types as, within 3D hydrogel systems, single-cell MSC encapsulation has been shown to match or outperform pellet/aggregate encapsulation [101], in contrast to the pellet culture method conventionally used to induce MSC chondrogenesis outside of a scaffold [102]. Thus, while both cell types were encapsulated using an identical single-cell suspension protocol, only iCPs spontaneously self-organized into multicellular aggregates. This divergence in aggregation behavior therefore reflects an intrinsic difference in developmental behavior between the two cell types rather than a difference in experimental handling, and highlights a practical advantage of the iCP system: unlike hMSCs, iCPs do not require an additional pre-formation step to form aggregates within non-functionalized alginate hydrogels, which may simplify future bioprocessing workflows. Interestingly, softer gels (1-2%) supported iCP aggregate formation whereas stiffer gels (3%) did not. These aggregates displayed a matrix composition rich in collagen type II and aggrecan, with an outer layer composed also of collagen type I, resembling cartilaginous pellet structures commonly used in cartilage disease modelling studies [39,74]. Within these aggregates, iCPs exhibited increased expression of hyaline cartilage-associated markers with reduced fibrocartilage (COL1) and hypertrophic (COL10) marker expression compared to hMSCs, with stiffness primarily influencing aggregates size and matrix deposition. This more hyaline-like phenotype of iCP-derived chondrocytes aligns with previous reports demonstrating the superior chondrogenic potential of chondroprogenitors over MSCs [26,31,33,41,42,73,103].

To investigate the behavior of MSCs and iCPs in hydrogels engineered to mimic more closely the native microenvironment, alginate gels were functionalized with developmentally relevant peptides. Given the established roles of integrin-mediated^109,110^ and cadherin-mediated^53,71,72,91^ signaling during cartilage formation, we investigated the effects of RGD and HAVDI peptide functionalization within 2% alginate gels. Functionalization (HAVDI/RGD and VAIDH/RGD) of 2% alginate gels did not significantly alter gel rheological properties but significantly altered cell behavior and matrix organization. For hMSCs, RGD functionalization increased collagen type I deposition while suppressing COL10 expression, suggesting these markers respond differently to integrin-mediated signaling. HAVDI incorporation significantly enhanced hMSCs hyaline marker expression without exacerbating hypertrophic gene expression, supporting previous findings on the beneficial effects of cadherin-mediated cell-cell interactions during chondrogenic differentiation [76–79,104]. However, its incorporation in this study did not mitigate collagen type I expression, as previously reported [76]. For iCPs, peptide-functionalization improved cellular distribution but only mildly increased viability. As noted above, the non-viable fraction likely represents off-target cells, and the selective loss is not detrimental: surviving cells consistently differentiated into chondrocytes, allowing day-12 iCPs to be used directly without additional purification, consistent with the original protocol’s pellet cultures, and avoiding a selection step that is typically both time-consuming and costly [40]. Functionalization, however, improved chondrogenesis of iCPs, though the response depended on peptide combination: RGD alone favored a more fibrocartilage-like profile, whereas HAVDI/RGD shifted expression toward a hyaline phenotype (SOX9, COL2, ACAN), suggesting that HAVDI incorporation offset the fibrocartilage-promoting effect observed with RGD alone. Moreover, RGD/HAVDI-functionalized gels exhibited uniform cellular distribution and ECM deposition, likely reflecting enhanced intercellular signaling, critical during early chondrogenesis [105]. iCPs also exhibited significantly lower COL10 expression than hMSCs in HAVDI/RGD gels, a difference accompanied by lower MMP13, an effector of hypertrophic matrix remodeling, across both peptide-functionalized conditions. RUNX2 and ALP, by contrast, did not differ between cell types, suggesting that the reduced hypertrophic markers expression in iCPs was not accompanied by a corresponding difference in upstream/mineralization-associated markers [106,107]. The uncoupling of RUNX2 from its downstream targets COL10 and MMP13 is consistent with reports that RUNX2 alone is insufficient to drive COL10a1 transcription and points to differences further downstream, such as BMP/Smad signaling [108] as a more likely explanation for the reduced hypertrophic profile of iCPs. Overall, our findings indicate that iCPs support a more hyaline-like chondrogenic phenotype than MSCs, particularly within biomaterials integrating both mechanical and bioactive cues.

While our findings demonstrate the excellent chondrogenic capacity of iCPs, several practical considerations merit discussion. This study employed a single MSC donor line and a single iPSC line, which limits assessment of inter-donor and inter-line variability. The iCP derivation protocol used in this study generates a heterogeneous population containing non-chondrogenic cells. Consequently, Wnt-C59 and ML329 inhibitors were required during differentiation to suppress off-target lineage commitment. This heterogeneity, combined with the lack of a validated iCP expansion protocol [33,39,73], requires the generation of fresh iCP batches for each experiment or careful cryopreservation strategies. In addition, iPSC maintenance and directed differentiation remain substantially more demanding in terms of cost, labor, and time than conventional MSCs culture [109]. However, these limitations are offset by several advantages: iPSCs can be readily expanded and genetically modified prior to differentiation, enabling patient-specific disease modelling and mechanistic studies of cartilage development through targeted gene editing [110]. Furthermore, continued improvements in iCP differentiation protocols [32] together with declining costs associated with iPSC technologies [110], may further support the broader adoption of iCP-based cartilage models. Nevertheless, further validation using multiple donor lines, long-term functional assessment will be required before translation toward regenerative or modelling applications.

## 5. CONCLUSIONS

This study compared MSC and iCP chondrogenesis within 3D alginate hydrogels and demonstrated the importance of tuning both gel mechanics and bioactive signaling cues to support chondrogenic differentiation. Among the tested conditions, soft, fast-relaxing alginate gels functionalized with both HAVDI and RGD peptides provided the most favorable environment for chondrogenic differentiation. In functionalized gels with HAVDI/RGD, iCP successfully developed into constructs that exhibited a more hyaline cartilage-like matrix composition and organization than those formed by hMSCs, supporting their use as an alternative cell source for cartilage tissue engineering and disease modelling. Future studies should aim to further elucidate the molecular mechanisms underlying differential cellular responses to biomaterial cues, with particular focus on the roles of integrins, cadherins, and mechanical signaling in regulating chondrogenic lineage commitment [96–99,111,112]. Additionally, incorporation of dynamic mechanical loading may further enhance ECM maturation and better recapitulate the biomechanical environment of native cartilage [113–115]. Together, these advances may support the establishment of more robust *in vitro* models of cartilage development, degeneration, and regenerative repair.

## Data Availability

The data that support the findings of this study are available from the corresponding author upon reasonable request.

## CRediT authorship contribution statement

**M.L.V.:** Conceptualization, Writing – original draft, review & editing, Visualization, Validation, Methodology, Investigation, Formal analysis, Data curation, Software. **A.R.T.:** Writing – review & editing, Supervision, Resources, Methodology. **M.J.D.:** Writing – review & editing, Supervision, Resources, Methodology. **N.K.**: Writing – review & editing, Supervision, Resources, Methodology. **C.A.:** Writing – review & editing. **B.D.**: Writing – review & editing, Methodology. **P.M.T.**: Writing – review & editing, Methodology, Resources. **T.S.**: Writing – review & editing, Validation. **C.H.**: Writing – review & editing, Supervision, Resources, Methodology, Conceptualization. **D.J.A.**: Writing – review & editing, Supervision, Resources, Project administration, Methodology, Funding acquisition, Conceptualization, Project administration. **C.S.G.**: Writing – review & editing, Supervision, Resources, Project administration, Methodology, Funding acquisition, Conceptualization.

## Funding sources

This work was supported by the SFI-EPSRC funded LifETIME Centre of Doctoral Training (EP/S02347X/1).

## Acknowledgements

The authors gratefully acknowledge the Flow Cytometry Core Lab and the Glasgow Imaging Facility at University of Glasgow for their support and assistance in this work, and REPROCELL Europe Ltd for their collaboration on this project. We are particularly grateful to David Bunton (CEO, REPROCELL Europe) for facilitating access to their facilities. We also thank Dr. Amanda Dicks, Zainab Harissa, and Dr Yu Seon Kim from the Department of Orthopaedic Surgery, Washington University School of Medicine in St. Louis, for their support with the chondroprogenitor differentiation protocol.

## Declaration of competing interest

REPROCELL Europe Ltd was a project stakeholder and N.K. was an employee of REPROCELL Europe Ltd during the study period. REPROCELL provided the iPSC cell line used in this study and iPSC culture training to the research team in their facilities. REPROCELL had no involvement in study design, data analysis, or manuscript preparation beyond N.K.’s contributions to iPSC culture training and manuscript editing. All other authors declare no competing interests.

## Notes

### Competing Interest Statement

Maria Laura Vieri reports equipment, drugs, or supplies were provided by REPROCELL Europe Ltd. Nikola Kolundzic reports a relationship with REPROCELL Europe Ltd that includes: employment and non-financial support. REPROCELL Europe Ltd was a project stakeholder and N.K. was an employee of REPROCELL Europe Ltd during the study period.
REPROCELL provided the iPSC cell line used in this study and iPSC culture training to the research team in their facilities. REPROCELL had no involvement in study design, data analysis, or manuscript preparation beyond N.K.'s contributions to iPSC culture training and manuscript editing. All other authors declare no competing interests.

## REFERENCES

[1] R. Wu, Y. Guo, Y. Chen, J. Zhang, Osteoarthritis burden and inequality from 1990 to 2021: a systematic analysis for the global burden of disease Study 2021, Sci. Rep. 15 (2025) 8305. 10.1038/s41598-025-93124-z.

[2] T. Stampoultzis, P. Karami, D.P. Pioletti, Thoughts on cartilage tissue engineering: A 21st century perspective, Curr. Res. Transl. Med. 69 (2021) 103299. 10.1016/j.retram.2021.103299.

[3] X. Guo, L. Xi, M. Yu, Z. Fan, W. Wang, A. Ju, Z. Liang, G. Zhou, W. Ren, Regeneration of articular cartilage defects: Therapeutic strategies and perspectives, J. Tissue Eng. 14 (2023) 20417314231164765. 10.1177/20417314231164765.

[4] H. Kwon, W.E. Brown, C.A. Lee, D. Wang, N. Paschos, J.C. Hu, K.A. Athanasiou, Surgical and tissue engineering strategies for articular cartilage and meniscus repair, Nat. Rev. Rheumatol. 15 (2019) 550–570. 10.1038/s41584-019-0255-1.

[5] M. Ansari, A. Darvishi, A. Sabzevari, A review of advanced hydrogels for cartilage tissue engineering, Front. Bioeng. Biotechnol. 12 (2024) 1340893. 10.3389/fbioe.2024.1340893.

[6] R.C. Nordberg, B.J. Bielajew, T. Takahashi, S. Dai, J.C. Hu, K.A. Athanasiou, Recent advancements in cartilage tissue engineering innovation and translation, Nat. Rev. Rheumatol. 20 (2024) 323–346. 10.1038/s41584-024-01118-4.

[7] P. Baei, H. Daemi, F. Aramesh, H. Baharvand, M.B. Eslaminejad, Advances in mechanically robust and biomimetic polysaccharide-based constructs for cartilage tissue engineering, Carbohydr. Polym. 308 (2023) 120650. 10.1016/j.carbpol.2023.120650.

[8] C. Ligorio, A. Mata, Synthetic extracellular matrices with function-encoding peptides, Nat. Rev. Bioeng. 1 (2023) 518–536. 10.1038/s44222-023-00055-3.

[9] T. Stampoultzis, P. Karami, D.P. Pioletti, Thoughts on cartilage tissue engineering: A 21st century perspective, Curr. Res. Transl. Med. 69 (2021) 103299. 10.1016/j.retram.2021.103299.

[10] R. Berebichez-Fridman, P.R. Montero-Olvera, Sources and Clinical Applications of Mesenchymal Stem Cells: State-of-the-art review, Sultan Qaboos Univ. Med. J. 18 (2018) e264–277. 10.18295/squmj.2018.18.03.002.

[11] A. Liras, Future research and therapeutic applications of human stem cells: general, regulatory, and bioethical aspects, J. Transl. Med. 8 (2010) 131. 10.1186/1479-5876-8-131.

[12] H. Le, W. Xu, X. Zhuang, F. Chang, Y. Wang, J. Ding, Mesenchymal stem cells for cartilage regeneration, J. Tissue Eng. 11 (2020) 2041731420943839. 10.1177/2041731420943839.

[13] N. Majumder, S. Ghosh, Unfolding the Mystery Behind the Onset of Chondrocyte Hypertrophy during Chondrogenesis: Toward Designing Advanced Permanent Cartilage-mimetic Biomaterials, Adv. Funct. Mater. 33 (2023) 2300651. 10.1002/adfm.202300651.

[14] C. Vinatier, J. Guicheux, Cartilage tissue engineering: From biomaterials and stem cells to osteoarthritis treatments, Ann. Phys. Rehabil. Med. 59 (2016) 139–144. 10.1016/j.rehab.2016.03.002.

[15] K. Zha, X. Li, Z. Yang, G. Tian, Z. Sun, X. Sui, Y. Dai, S. Liu, Q. Guo, Heterogeneity of mesenchymal stem cells in cartilage regeneration: from characterization to application, Npj Regen. Med. 6 (2021) 14. 10.1038/s41536-021-00122-6.

[16] J.J. Bara, R.G. Richards, M. Alini, M.J. Stoddart, Concise Review: Bone Marrow-Derived Mesenchymal Stem Cells Change Phenotype Following In Vitro Culture: Implications for Basic Research and the Clinic, Stem Cells 32 (2014) 1713–1723. 10.1002/stem.1649.

[17] J. Phelps, A. Sanati-Nezhad, M. Ungrin, N.A. Duncan, A. Sen, Bioprocessing of Mesenchymal Stem Cells and Their Derivatives: Toward Cell-Free Therapeutics, Stem Cells Int. 2018 (2018) 9415367. 10.1155/2018/9415367.

[18] W. (first) Tsuji, P.J. Rubin, K.G. Marra, Adipose-derived stem cells: Implications in tissue regeneration, World J. Stem Cells 6 (2014) 312. 10.4252/wjsc.v6.i3.312.

[29] K. Miclau, W.S. Hambright, J. Huard, M.J. Stoddart, C.S. Bahney, Cellular expansion of MSCs: Shifting the regenerative potential, Aging Cell 22 (2023) e13759. 10.1111/acel.13759.

[20] R.M. Alves-Paiva, S. do Nascimento, D. De Oliveira, L. Coa, K. Alvarez, N. Hamerschlak, O.K. Okamoto, L.C. Marti, A.T. Kondo, J.M. Kutner, M.A.T. Bortolini, R. Castro, J.A.P. de Godoy, Senescence State in Mesenchymal Stem Cells at Low Passages: Implications in Clinical Use, Front. Cell Dev. Biol. 10 (2022). https://www.frontiersin.org/articles/10.3389/fcell.2022.858996.

[21] C. Shigley, J. Trivedi, O. Meghani, B.D. Owens, C.T. Jayasuriya, Suppressing Chondrocyte Hypertrophy to Build Better Cartilage, Bioengineering 10 (2023). 10.3390/bioengineering10060741.

[22] R.A. Somoza, J.F. Welter, D. Correa, A.I. Caplan, Chondrogenic differentiation of mesenchymal stem cells: challenges and unfulfilled expectations, Tissue Eng. Part B Rev. 20 (2014) 596–608. 10.1089/ten.TEB.2013.0771.

[23] E. Steck, H. Bertram, R. Abel, B. Chen, A. Winter, W. Richter, Induction of Intervertebral Disc–Like Cells From Adult Mesenchymal Ste m Cells, Stem Cells 23 (2005) 403–411. 10.1634/stemcells.2004-0107.

[24] J.C. Bernhard, G. Vunjak-Novakovic, Should we use cells, biomaterials, or tissue engineering for cartilage regeneration?, Stem Cell Res. Ther. 7 (2016) 56. 10.1186/s13287-016-0314-3.

[25] E. Vinod, K. Padmaja, A. Livingston, J.V. James, S.M. Amirtham, S. Sathishkumar, B. Ramasamy, G. Rebekah, A.J. Daniel, U. Kachroo, Prospective Isolation and Characterization of Chondroprogenitors from Human Chondrocytes Based on CD166/CD34/CD146 Surface Markers, Cartilage 13 (2021). 10.1177/19476035211042412.

[26] E. Vinod, R. Parameswaran, B. Ramasamy, U. Kachroo, Pondering the Potential of Hyaline Cartilage–Derived Chondroprogenitors for Tissue Regeneration: A Systematic Review, Cartilage 13 (2021) 34S–52S. 10.1177/1947603520951631.

[27] M. Rikkers, J.V. Korpershoek, R. Levato, J. Malda, L.A. Vonk, The clinical potential of articular cartilage-derived progenitor cells: a systematic review, Npj Regen. Med. 7 (2022) 2. 10.1038/s41536-021-00203-6.

[28] R. Levato, W.R. Webb, I.A. Otto, A. Mensinga, Y. Zhang, M. Rijen, R. Weeren, I.M. Khan, J. Malda, The bio in the ink: cartilage regeneration with bioprintable hydrogels and articular cartilage-derived progenitor cells, Acta Biomater. 61 (2017) 41–53. 10.1016/j.actbio.2017.08.005.

[29] N. Nakayama, A. Pothiawala, J.Y. Lee, N. Matthias, K. Umeda, B.K. Ang, J. Huard, Y. Huang, D. Sun, Human pluripotent stem cell-derived chondroprogenitors for cartilage tissue engineering, Cell. Mol. Life Sci. 77 (2020) 2543–2563. 10.1007/s00018-019-03445-2.

[30] C. Feng, W.C.W. Chan, Y. Lam, X. Wang, P. Chen, B. Niu, V.C.W. Ng, J.C. Yeo, S. Stricker, K.S.E. Cheah, M. Koch, S. Mundlos, H.H. Ng, D. Chan, Lgr5 and Col22a1 Mark Progenitor Cells in the Lineage toward Juvenile Articular Chondrocytes, Stem Cell Rep. 13 (2019) 713–729. 10.1016/j.stemcr.2019.08.006.

[31] C.T. Jayasuriya, Q. Chen, Potential benefits and limitations of utilizing chondroprogenitors in cell-based cartilage therapy, Connect. Tissue Res. 56 (2015) 265–271. 10.3109/03008207.2015.1040547.

[32] P.A. Humphreys, F.E. Mancini, M.J.S. Ferreira, S. Woods, L. Ogene, S.J. Kimber, Developmental principles informing human pluripotent stem cell differentiation to cartilage and bone, Semin. Cell Dev. Biol. 127 (2022) 17–36. 10.1016/j.semcdb.2021.11.024.

[33] A.R. Dicks, N. Steward, F. Guilak, C.-L. Wu, Chondrogenic Differentiation of Human-Induced Pluripotent Stem Cells, in: M.J. Stoddart, E. Della Bella, A.R. Armiento (Eds.), Cartil. Tissue Eng., Springer US, New York, NY, 2023: pp. 87–114. 10.1007/978-1-0716-2839-3_8.

[34] A.Y. Owaidah, Induced pluripotent stem cells in cartilage tissue engineering: a literature review, Biosci. Rep. 44 (2024) BSR20232102. 10.1042/BSR20232102.

[35] K. Takahashi, S. Yamanaka, Induction of Pluripotent Stem Cells from Mouse Embryonic and Adult Fibroblast Cultures by Defined Factors, Cell 126 (2006) 663–676. 10.1016/j.cell.2006.07.024.

[36] J.J. Hwang, J. Choi, Y.A. Rim, Y. Nam, J.H. Ju, Application of Induced Pluripotent Stem Cells for Disease Modeling and 3D Model Construction: Focus on Osteoarthritis, Cells 10 (2021). 10.3390/cells10113032.

[37] C. Liu, A. Oikonomopoulos, N. Sayed, J.C. Wu, Modeling human diseases with induced pluripotent stem cells: from 2D to 3D and beyond, Development 145 (2018) dev156166. 10.1242/dev.156166.

[38] B.S. Ludwig, H. Kessler, S. Kossatz, U. Reuning, RGD-Binding Integrins Revisited: How Recently Discovered Functions and Novel Synthetic Ligands (Re-)Shape an Ever-Evolving Field, Cancers 13 (2021). 10.3390/cancers13071711.

[39] C.-L. Wu, A. Dicks, N. Steward, R. Tang, D.B. Katz, Y.-R. Choi, F. Guilak, Single cell transcriptomic analysis of human pluripotent stem cell chondrogenesis, Nat. Commun. 12 (2021) 362. 10.1038/s41467-020-20598-y.

[40] A. Dicks, C.-L. Wu, N. Steward, S.S. Adkar, C.A. Gersbach, F. Guilak, Prospective isolation of chondroprogenitors from human iPSCs based on cell surface markers identified using a CRISPR-Cas9-generated reporter, Stem Cell Res. Ther. 11 (2020) 66. 10.1186/s13287-020-01597-8.

[41] A. Pothiawala, B.E. Sahbazoglu, B.K. Ang, N. Matthias, G. Pei, Q. Yan, B.R. Davis, J. Huard, Z. Zhao, N. Nakayama, GDF5+ chondroprogenitors derived from human pluripotent stem cells preferentially form permanent chondrocytes, Development 149 (2022) dev196220. 10.1242/dev.196220.

[42] A. Rodríguez Ruiz, A. Dicks, M. Tuerlings, K. Schepers, M. Pel, R.G.H.H. Nelissen, C. Freund, C.L. Mummery, V. Orlova, F. Guilak, I. Meulenbelt, Y.F.M. Ramos, Cartilage from human-induced pluripotent stem cells: comparison with neo-cartilage from chondrocytes and bone marrow mesenchymal stromal cel ls, Cell Tissue Res. 386 (2021) 309–320. 10.1007/s00441-021-03498-5.

[43] L. Li, F. Yu, L. Zheng, R. Wang, W. Yan, Z. Wang, J. Xu, J. Wu, D. Shi, L. Zhu, X. Wang, Q. Jiang, Natural hydrogels for cartilage regeneration: Modification, preparation and application, J. Orthop. Transl. 17 (2019) 26–41. 10.1016/j.jot.2018.09.003.

[44] W. Liu, H. Madry, M. Cucchiarini, Application of Alginate Hydrogels for Next-Generation Articular Cartilage Regeneration, Int. J. Mol. Sci. 23 (2022) 1147. 10.3390/ijms23031147.

[45] B. Bachmann, S. Spitz, B. Schädl, A.H. Teuschl, H. Redl, S. Nürnberger, P. Ertl, Stiffness Matters: Fine-Tuned Hydrogel Elasticity Alters Chondrogenic Redifferentiation, Front. Bioeng. Biotechnol. (2020). 10.3389/fbioe.2020.00373.

[46] A.J. Engler, S. Sen, H.L. Sweeney, D.E. Discher, Matrix Elasticity Directs Stem Cell Lineage Specification, Cell 126 (2006) 677–689. 10.1016/j.cell.2006.06.044.

[47] O. Chaudhuri, Viscoelastic hydrogels for 3D cell culture, Biomater. Sci. 5 (2017) 1480– 1490. 10.1039/C7BM00261K.

[48] M. Walker, E.W. Pringle, G. Ciccone, L. Oliver-Cervelló, M. Tassieri, D. Gourdon, M. Cantini, Mind the Viscous Modulus: The Mechanotransductive Response to the Viscous Nature of Isoelastic Matrices Regulates Stem Cell Chondrogenesis, Adv. Healthc. Mater. n/a (2023) 2302571. 10.1002/adhm.202302571.

[49] S.L. Vega, M.Y. Kwon, K.H. Song, C. Wang, R.L. Mauck, L. Han, J.A. Burdick, Combinatorial hydrogels with biochemical gradients for screening 3D cellular microenvironments, Nat. Commun. 9 (2018) 614. 10.1038/s41467-018-03021-5.

[50] M.P. Dieterle, A. Husari, B. Rolauffs, T. Steinberg, P. Tomakidi, Integrins, cadherins and channels in cartilage mechanotransduction: perspectives for future regeneration strategies, Expert Rev. Mol. Med. 23 (2021) e14. 10.1017/erm.2021.16.

[51] W. Ke, L. Ma, B. Wang, Y. Song, R. Luo, G. Li, Z. Liao, Y. Shi, K. Wang, X. Feng, S. Li, W. Hua, C. Yang, N-cadherin mimetic hydrogel enhances MSC chondrogenesis through cell metabolism, Acta Biomater. 150 (2022) 83–95. 10.1016/j.actbio.2022.07.050.

[52] Y. Wang, Y. Xiao, S. Long, Y. Fan, X. Zhang, Role of N-Cadherin in a Niche-Mimicking Microenvironment for Chondrogenesis of Mesenchymal Stem Cells In Vitro, ACS Biomater. Sci. Eng. 6 (2020) 3491–3501. 10.1021/acsbiomaterials.0c00149.

[53] J. Dumbleton, P. Agarwal, H. Huang, N. Hogrebe, R. Han, K.J. Gooch, X. He, The Effect of RGD Peptide on 2D and Miniaturized 3D Culture of HEPM Cells, MSCs, and ADSCs with Alginate Hydrogel, Cell. Mol. Bioeng. 9 (2016) 277–288. 10.1007/s12195-016-0428-9.

[54] P. Singh, J.E. Schwarzbauer, Fibronectin and stem cell differentiation – lessons from chondrogenesis, J. Cell Sci. 125 (2012) 3703–3712. 10.1242/jcs.095786.

[55] P. Sharma, J.D. Twomey, M. Patkin, A.H. Hsieh, Layered alginate constructs: A platform for co-culture of heterogeneous cell populations, J. Vis. Exp. 2016 (2016) 1–6. 10.3791/54380.

[56] G. Ciccone, O. Dobre, G.M. Gibson, J.M. Rey, C. Gonzalez–Garcia, M. Vassalli, M. Salmeron–Sanchez, M. Tassieri, What Caging Force Cells Feel in 3D Hydrogels: A Rheological Perspective, Adv. Healthc. Mater. 9 (2020) 2000517. 10.1002/adhm.202000517.

[57] B. Henkel, T. Metzenthin, M. Gerken, W. Fiedler, Capping of unprotected amino groups during peptide synthesis, US11028123B2, 2021. https://patents.google.com/patent/US11028123B2/en (accessed May 13, 2026).

[58] J.A. Rowley, G. Madlambayan, D.J. Mooney, Alginate hydrogels as synthetic extracellular matrix materials, Biomaterials 20 (1999) 45–53. 10.1016/S0142-9612(98)00107-0.

[59] T. Hodgkinson, P.M. Tsimbouri, V. Llopis-Hernandez, P. Campsie, D. Scurr, P.G. Childs, D. Phillips, S. Donnelly, J.A. Wells, F.J. O’Brien, M. Salmeron-Sanchez, K. Burgess, M. Alexander, M. Vassalli, R.O.C. Oreffo, S. Reid, D.J. France, M.J. Dalby, The use of nanovibration to discover specific and potent bioactive metabolites that stimulate osteogenic differentiation in mesenchymal stem cells, Sci. Adv. 7 (2021) eabb7921. 10.1126/sciadv.abb7921.

[60] A.X. Sun, H. Lin, M.R. Fritch, H. Shen, P.G. Alexander, M. DeHart, R.S. Tuan, Chondrogenesis of human bone marrow mesenchymal stem cells in 3-dimensional, photocrosslinked hydrogel constructs: Effect of cell seeding density and material stiffness, Acta Biomater. 58 (2017) 302–311. 10.1016/j.actbio.2017.06.016.

[61] M. Walker, J. Luo, E.W. Pringle, M. Cantini, ChondroGELesis: Hydrogels to harness the chondrogenic potential of stem cells, Mater. Sci. Eng. C 121 (2021) 111822. 10.1016/j.msec.2020.111822.

[62] S. Lin, N. Sangaj, T. Razafiarison, C. Zhang, S. Varghese, Influence of Physical Properties of Biomaterials on Cellular Behavior, Pharm. Res. 28 (2011) 1422–1430. 10.1007/s11095-011-0378-9.

[63] H. Lee, L. Gu, D.J. Mooney, M.E. Levenston, O. Chaudhuri, Mechanical confinement regulates cartilage matrix formation by chondrocytes, Nat. Mater. 16 (2017) 1243– 1251. 10.1038/nmat4993.

[64] H. Ying, C. Shen, R. Pan, X. Li, Y. Chen, Strategy insight: Mechanical properties of biomaterials’ influence on hydrogel-mesenchymal stromal cell combination for osteoarthritis therapy, Front. Pharmacol. Volume 14–2023 (2023). https://www.frontiersin.org/journals/pharmacology/articles/10.3389/fphar.2023.1152612.

[65] A.D. Augst, H.J. Kong, D.J. Mooney, Alginate Hydrogels as Biomaterials, Macromol. Biosci. 6 (2006) 623–633. 10.1002/mabi.200600069.

[66] J. Zhang, E. Wehrle, J.R. Vetsch, G.R. Paul, M. Rubert, R. Müller, Alginate dependent changes of physical properties in 3D bioprinted cell-laden porous scaffolds affect cell viability and cell morphology, Biomed. Mater. 14 (2019) 065009. 10.1088/1748-605X/ab3c74.

[67] O. Chaudhuri, L. Gu, D. Klumpers, M. Darnell, S.A. Bencherif, J.C. Weaver, N. Huebsch, H.P. Lee, E. Lippens, G.N. Duda, D.J. Mooney, Hydrogels with tunable stress relaxation regulate stem cell fate and activity, Nat. Mater. 15 (2016) 326–334. 10.1038/nmat4489.

[68] M.M.J. Caron, P.J. Emans, M.M.E. Coolsen, L. Voss, D.A.M. Surtel, A. Cremers, L.W. van Rhijn, T.J.M. Welting, Redifferentiation of dedifferentiated human articular chondrocytes: comparison of 2D and 3D cultures, Osteoarthritis Cartilage 20 (2012) 1170–1178. 10.1016/j.joca.2012.06.016.

[69] Y. Chen, Y. Yu, Y. Wen, J. Chen, J. Lin, Z. Sheng, W. Zhou, H. Sun, C. An, J. Chen, W. Wu, C. Teng, W. Wei, H. Ouyang, A high-resolution route map reveals distinct stages of chondrocyte dedifferentiation for cartilage regeneration, Bone Res. 10 (2022) 38. 10.1038/s41413-022-00209-w.

[70] X. Zhou, K. von der Mark, S. Henry, W. Norton, H. Adams, B. de Crombrugghe, Chondrocytes Transdifferentiate into Osteoblasts in Endochondral Bone during Development, Postnatal Growth and Fracture Healing in Mice, PLOS Genet. 10 (2014) e1004820. 10.1371/journal.pgen.1004820.

[71] E. Charlier, C. Deroyer, F. Ciregia, O. Malaise, S. Neuville, Z. Plener, M. Malaise, D. de Seny, Chondrocyte dedifferentiation and osteoarthritis (OA), Biochem. Pharmacol. 165 (2019) 49–65. 10.1016/j.bcp.2019.02.036.

[72] P.G. Bush, A.C. Hall, The volume and morphology of chondrocytes within non-degenerate and degenerate human articular cartilage, Osteoarthritis Cartilage 11 (2003) 242–251. 10.1016/S1063-4584(02)00369-2.

[73] S.S. Adkar, C.-L. Wu, V.P. Willard, A. Dicks, A. Ettyreddy, N. Steward, N. Bhutani, C.A. Gersbach, F. Guilak, Step-Wise Chondrogenesis of Human Induced Pluripotent Stem Cells and Purification Via a Reporter Allele Generated by CRISPR-Cas9 Genome Editing, Stem Cells 37 (2019) 65–76. 10.1002/stem.2931.

[74] A.R. Dicks, G.I. Maksaev, Z. Harissa, A. Savadipour, R. Tang, N. Steward, W. Liedtke, C.G. Nichols, C.-L. Wu, F. Guilak, Skeletal dysplasia-causing TRPV4 mutations suppress the hypertrophic differentiation of human iPSC-derived chondrocytes, eLife 12 (2023) e71154. 10.7554/eLife.71154.

[75] M.E. Candela, R. Yasuhara, M. Iwamoto, M. Enomoto-Iwamoto, Resident mesenchymal progenitors of articular cartilage, Matrix Biol. 39 (2014) 44–49. 10.1016/j.matbio.2014.08.015.

[76] L. Bian, M. Guvendiren, R.L. Mauck, J.A. Burdick, Hydrogels that mimic developmentally relevant matrix and N-cadherin interactions enhance MSC chondrogenesis, Proc. Natl. Acad. Sci. 110 (2013) 10117–10122. 10.1073/pnas.1214100110.

[77] M.Y. Kwon, S.L. Vega, W.M. Gramlich, M. Kim, R.L. Mauck, J.A. Burdick, Dose and Timing of N-Cadherin Mimetic Peptides Regulate MSC Chondrogenesis within Hydrogels, Adv. Healthc. Mater. 7 (2018) 1–10. 10.1002/adhm.201701199.

[78] T.H. Qazi, D.J. Mooney, G.N. Duda, S. Geissler, Niche-mimicking interactions in peptide-functionalized 3D hydrogels amplify mesenchymal stromal cell paracrine effects, Biomaterials 230 (2020) 119639. 10.1016/j.biomaterials.2019.119639.

[79] Z. Zhang, B. Sha, L. Zhao, H. Zhang, J. Feng, C. Zhang, L. Sun, M. Luo, B. Gao, H. Guo, Z. Wang, F. Xu, T.J. Lu, G.M. Genin, M. Lin, Programmable integrin and N-cadherin adhesive interactions modulate mechanosensing of mesenchymal stem cells by cofilin phosphorylation, Nat. Commun. 13 (2022) 6854. 10.1038/s41467-022-34424-0.

[80] W.M. Kulyk, W.B. Upholt, R.A. Kosher, Fibronectin gene expression during limb cartilage differentiation, Development 106 (1989) 449–455. 10.1242/dev.106.3.449.

[81] P. Singh, J.E. Schwarzbauer, Fibronectin and stem cell differentiation – lessons from chondrogenesi s, J. Cell Sci. 125 (2012) 3703–3712. 10.1242/jcs.095786.

[82] A.R. Tan, C.T. Hung, Concise Review: Mesenchymal Stem Cells for Functional Cartilage Tissue Engineering: Taking Cues from Chondrocyte-Based Constructs, Stem Cells Transl. Med. 6 (2017) 1295–1303. 10.1002/sctm.16-0271.

[83] C. Manferdini, D. Trucco, Y. Saleh, E. Gabusi, P. Dolzani, E. Lenzi, L. Vannozzi, L. Ricotti, G. Lisignoli, RGD-Functionalized Hydrogel Supports the Chondrogenic Commitment of Adipose Mesenchymal Stromal Cells, Gels 8 (2022). 10.3390/gels8060382.

[84] J.T. Connelly, A.J. García, M.E. Levenston, Inhibition of in vitro chondrogenesis in RGD-modified three-dimensional alginate gels, Biomaterials 28 (2007) 1071–1083. 10.1016/j.biomaterials.2006.10.006.

[85] T. Re’em, O. Tsur-Gang, S. Cohen, The effect of immobilized RGD peptide in macroporous alginate scaffolds on TGFβ1-induced chondrogenesis of human mesenchymal stem cells, Biomaterials 31 (2010) 6746–6755. 10.1016/j.biomaterials.2010.05.025.

[86] A.K. Kudva, F.P. Luyten, J. Patterson, RGD-functionalized polyethylene glycol hydrogels support proliferation and in vitro chondrogenesis of human periosteum-derived cells, J. Biomed. Mater. Res. A 106 (2018) 33–42. 10.1002/jbm.a.36208.

[87] T.H. Qazi, D.J. Mooney, G.N. Duda, S. Geissler, Niche-mimicking interactions in peptide-functionalized 3D hydrogels am plify mesenchymal stromal cell paracrine effects, Biomaterials 230 (2020) 119639. 10.1016/j.biomaterials.2019.119639.

[88] Z. Zhang, B. Sha, L. Zhao, H. Zhang, J. Feng, C. Zhang, L. Sun, M. Luo, B. Gao, H. Guo, Z. Wang, F. Xu, T.J. Lu, G.M. Genin, M. Lin, Programmable integrin and N-cadherin adhesive interactions modulate me chanosensing of mesenchymal stem cells by cofilin phosphorylation, Nat. Commun. 13 (2022) 6854. 10.1038/s41467-022-34424-0.

[89] Z. Ge, Y. Hu, B.C. Heng, Z. Yang, H. Ouyang, E.H. Lee, T. Cao, Osteoarthritis and therapy, Arthritis Care Res. 55 (2006) 493–500. 10.1002/art.21994.

[90] D. Huang, Y. Li, Z. Ma, H. Lin, X. Zhu, Y. Xiao, X. Zhang, Collagen hydrogel viscoelasticity regulates MSC chondrogenesis in a RO CK-dependent manner, Sci. Adv. 9 (n.d.) eade9497. 10.1126/sciadv.ade9497.

[91] A.J. Steward, D.J. Kelly, Mechanical regulation of mesenchymal stem cell differentiation, J Anat 227 (2015) 717–731. 10.1111/joa.12243.

[92] A. Karim, A.K. Amin, A.C. Hall, The clustering and morphology of chondrocytes in normal and mildly degenerate human femoral head cartilage studied by confocal laser scanning microscopy, J Anat 232 (2018) 686–698. 10.1111/joa.12768.

[93] W. Yan, M. Maimaitimin, Y. Wu, Y. Fan, S. Ren, F. Zhao, C. Cao, X. Hu, J. Cheng, Y. Ao, Meniscal fibrocartilage regeneration inspired by meniscal maturational and regenerative process, Sci. Adv. 9 (2023) eadg8138. 10.1126/sciadv.adg8138.

[94] W.-T. Yan, J.-S. Wang, P.-Z. Fan, S. Roberts, K. Wright, Z.-Z. Zhang, The clinical potential of meniscal progenitor cells, J. Cartil. Jt. Preserv. 4 (2024) 100166. 10.1016/j.jcjp.2024.100166.

[95] T. Jörimann, P. Füllemann, A. Jose, R. Matthys, E. Wehrle, M.J. Stoddart, S. Verrier, In Vitro Induction of Hypertrophic Chondrocyte Differentiation of Naïve MSCs by Strain, Cells 14 (2025) 25. 10.3390/cells14010025.

[96] B. Shu, M. Zhang, R. Xie, M. Wang, H. Jin, W. Hou, D. Tang, S.E. Harris, Y. Mishina, R.J. O’Keefe, M.J. Hilton, Y. Wang, D. Chen, BMP2, but not BMP4, is crucial for chondrocyte proliferation and maturation during endochondral bone development, J. Cell Sci. 124 (2011) 3428–3440. 10.1242/jcs.083659.

[97] A.M. DeLise, L. Fischer, R.S. Tuan, Cellular interactions and signaling in cartilage development, Osteoarthritis Cartilage 8 (2000) 309–334. 10.1053/joca.1999.0306.

[98] A.M. DeLise, R.S. Tuan, Alterations in the spatiotemporal expression pattern and function of N-Cadherin inhibit cellular condensation and chondrogenesis of limb mesenchymal cells in vitro, J. Cell. Biochem. 87 (2002) 342–359. 10.1002/jcb.10308.

[99] H.M. Kronenberg, Developmental regulation of the growth plate, Nature 423 (2003) 332–336. 10.1038/nature01657.

[100] S. Ghosh, M. Laha, S. Mandal, S. Sengupta, D.L. Kaplan, In vitro Model of Mesenchymal Condensation During Chondrogenic Development, Biomaterials 30 (2009) 6530–6540. 10.1016/j.biomaterials.2009.08.019.

[101] H. Rogan, F. Ilagan, F. Yang, Comparing Single Cell Versus Pellet Encapsulation of Mesenchymal Stem Cells in Three-Dimensional Hydrogels for Cartilage Regeneration, Tissue Eng. Part A 25 (2019) 1404–1412. 10.1089/ten.tea.2018.0289.

[102] B. Johnstone, T.M. Hering, A.I. Caplan, V.M. Goldberg, J.U. Yoo, *In Vitro*Chondrogenesis of Bone Marrow-Derived Mesenchymal Progenitor Cells, Exp. Cell Res. 238 (1998) 265–272. 10.1006/excr.1997.3858.

[103] E. Vinod, G. Parasuraman, J. Lisha J., S.M. Amirtham, A. Livingston, J.J. Varghese, S. Rani, D.V. Francis, G. Rebekah, A.J. Daniel, B. Ramasamy, S. Sathishkumar, Human fetal cartilage-derived chondrocytes and chondroprogenitors display a greater commitment to chondrogenesis than adult cartilage resident cells, PLoS ONE 18 (2023) e0285106. 10.1371/journal.pone.0285106.

[104] Y. Rong, Z. Zhang, C. He, X. Chen, Bioactive polypeptide hydrogels modified with RGD and N-cadherin mimetic peptide promote chondrogenic differentiation of bone marrow mesenchymal stem cells, Sci. China Chem. 63 (2020) 1100–1111. 10.1007/s11426-020-9772-0.

[105] R. Li, J. Xu, D.S.H. Wong, J. Li, P. Zhao, L. Bian, Self-assembled N-cadherin mimetic peptide hydrogels promote the chondrogenesis of mesenchymal stem cells through inhibition of canonical Wnt/β-catenin signaling, Biomaterials 145 (2017) 33–43. 10.1016/j.biomaterials.2017.08.031.

[106] M. D’Angelo, Z. Yan, M. Nooreyazdan, M. Pacifici, D. s. Sarment, P. c. Billings, P. s. Leboy, MMP-13 is induced during chondrocyte hypertrophy, J. Cell. Biochem. 77 (2000) 678–693. 10.1002/(SICI)1097-4644(20000615)77:4%3C678::AID-JCB15%3E3.0.CO;2-P.

[107] D. Miao, A. Scutt, Histochemical Localization of Alkaline Phosphatase Activity in Decalcified Bone and Cartilage, J. Histochem. Cytochem. 50 (2002) 333–340. 10.1177/002215540205000305.

[108] J. Gu, Y. Lu, F. Li, L. Qiao, Q. Wang, N. Li, J.A. Borgia, Y. Deng, G. Lei, Q. Zheng, Identification and characterization of the novel Col10a1 regulatory mechanism during chondrocyte hypertrophic differentiation, Cell Death Dis. 5 (2014) e1469–e1469. 10.1038/cddis.2014.444.

[109] M. Madrid, U. Lakshmipathy, X. Zhang, K. Bharti, D.M. Wall, Y. Sato, G. Muschler, A. Ting, N. Smith, S. Deguchi, S. Kawamata, J.C. Moore, B. Makovoz, S. Sullivan, V. Falco, A.Z. Al-Riyami, Considerations for the development of iPSC-derived cell therapies: a review of key challenges by the JSRM-ISCT iPSC Committee, Cytotherapy 26 (2024) 1382–1399. 10.1016/j.jcyt.2024.05.022.

[110] P. De Kinderen, J. Meester, B. Loeys, S. Peeters, E. Gouze, S. Woods, G. Mortier, A. Verstraeten, Differentiation of Induced Pluripotent Stem Cells Into Chondrocytes: Methods and Applications for Disease Modeling and Drug Discovery, J. Bone Miner. Res. Off. J. Am. Soc. Bone Miner. Res. 37 (2022) 397–410. 10.1002/jbmr.4524.

[111] R.F. Loeser, Integrins and chondrocyte–matrix interactions in articular cartilage, Matrix Biol. 39 (2014) 11–16. 10.1016/j.matbio.2014.08.007.

[112] M.P. Dieterle, A. Husari, B. Rolauffs, T. Steinberg, P. Tomakidi, Integrins, cadherins and channels in cartilage mechanotransduction: pe rspectives for future regeneration strategies, Expert Rev. Mol. Med. 23 (2021) e14. 10.1017/erm.2021.16.

[113] A. Abdal Dayem, A. Prince, A.M.M. Gabr, Chondrogenic Differentiation of Stem Cells for Cartilage Regeneration: Advances and Future Perspectives, Tissue Eng. Regen. Med. 23 (2026) 21–84. 10.1007/s13770-025-00768-z.

[114] L. Mecchi, M.M.J. Caron, T.J.M. Welting, M.J. Stoddart, The impact of mechanical bioreactors on human mesenchymal stromal cells utilized for articular cartilage repair, Acta Biomater. 210 (2026) 40–56. 10.1016/j.actbio.2025.12.029.

[115] J.R. Choi, K.W. Yong, J.Y. Choi, Effects of mechanical loading on human mesenchymal stem cells for cartilage tissue engineering, J. Cell. Physiol. 233 (2018) 1913–1928. 10.1002/jcp.26018.

