## Supplementary Figures for "iPSC-Derived Chondroprogenitors as a Promising Cell Source for Cartilage Engineering: A Comparison with MSCs in Unmodified and Peptide-Functionalized Alginate Hydrogels"

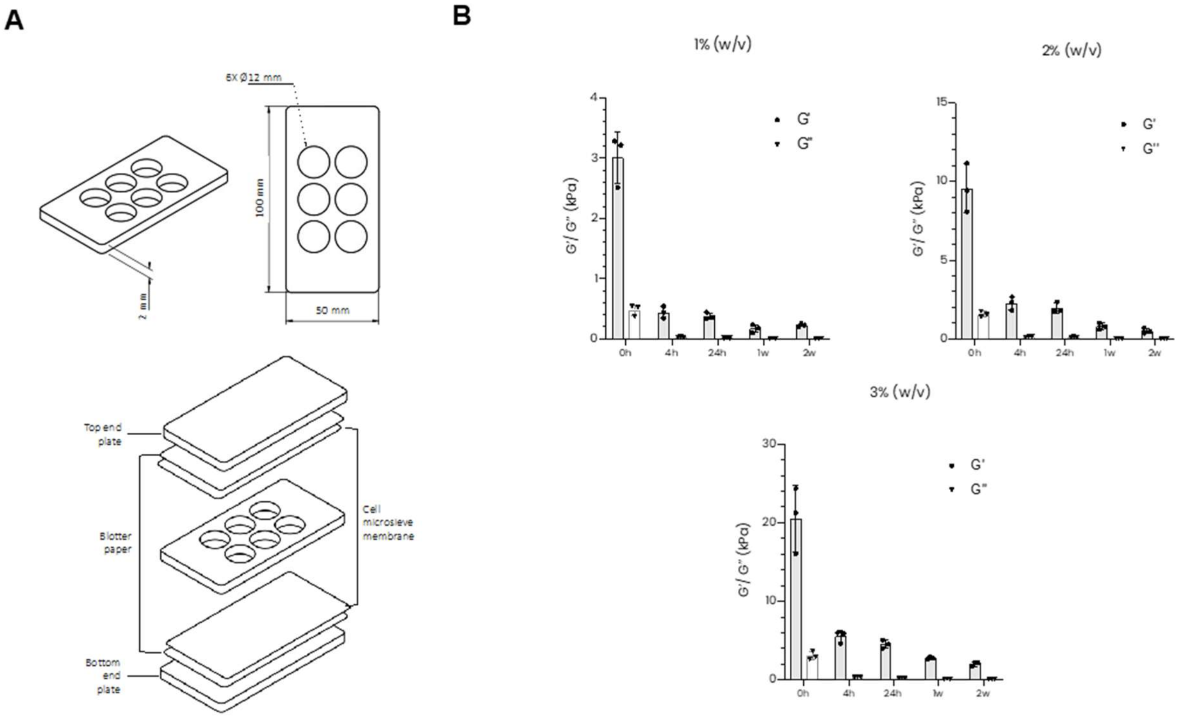


**Figure S1 | A)** Dimensions of the plastic molds used for fabrication of gel discs (top) and schematic representation of mold assembly (bottom). The central welled component is enclosed between two cell microsieve membranes, two blotter papers, and the end plates. **B)** Amplitude sweep analysis of 1-3 % (w/v) alginate hydrogels in cell culture medium, measuring storage modulus (G′) and loss modulus (G″) over time. All formulations maintained gel-like behavior (G′> G″) at all time points. Error bars represent mean ± s.d. (n = 3).

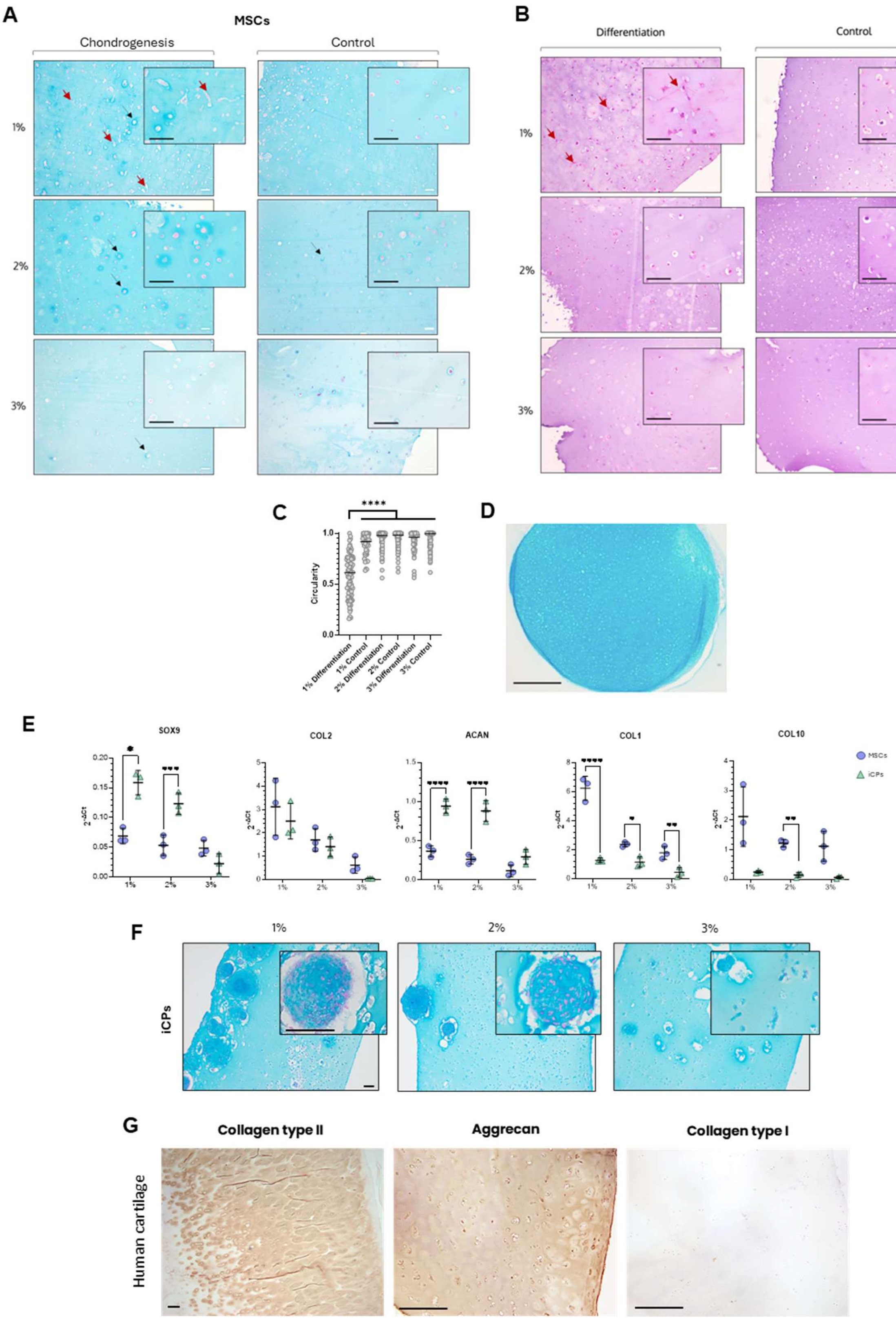
A
MSCs
Chondrogenesis
Control
1%
2%
3%
B
Differentiation
Control
1%
2%
3%
C
****
Circularity
1.0
0.5
0.0
1% Differentiation
1% Control
2% Differentiation
2% Control
3% Differentiation
3% Control
D
E
SOX9
COL2
ACAN
COL1
COL10
MSCs
iCPs
2-ΔCt
1%
2%
3%
F
1%
2%
3%
iCPs
G
Collagen type II
Aggrecan
Collagen type I
Human cartilage

**Figure S2 | Histological analysis of 1-3% (w/v) unmodified alginate gels. A)** Alcian blue and hematoxylin staining of paraffin-embedded sections of hMSCs cultured in 1-3% (w/v) unmodified gels after 28 days in chondrogenic and control medium. Due to the negatively charged nature of alginate, hydrogels stain blue; however, proteoglycan deposition is observed as a more intense blue halo surrounding the cells (examples indicated by black arrows). Elongated cell morphology is indicated by red arrows. Experiments were repeated three times with comparable results. Scale bar = 100 µm. **B)** Hematoxylin and eosin (H&E) staining of hMSCs embedded in 1-3% (w/v) unmodified gels after 28 days in chondrogenic or control medium. Elongated cells are indicated by red arrows. Scale bars = 100 µm. **C)** Quantification of cell circularity of hMSCs cultured in 1–3% (w/v) unmodified gels. One-way ANOVA with Tukey's multiple comparisons post-hoc test was performed (n = 48-87 from two biological replicates). **D)** Representative Alcian blue staining of iCPs pellets at day 28. Scale bars = 100 µm. **E)** Gene expression comparison between hMSCs and iCPs in 1-3% (w/v) gels after 28 days of chondrogenesis. Statistical analysis was performed using multiple t-test with Holm-Šídák correction for multiple comparisons. Error bars represent mean ± s.d. (n = 3). **F)** Alcian blue and hematoxylin staining of paraffin-embedded sections of iCPs cultured in 1-3% alginate gels after 28 days under chondrogenic conditions. Experiments were repeated three times with comparable results. Scale bar = 100 µm. **G)** Positive controls for IHC: sections from healthy regions of osteoarthritic human cartilage were used to validate antibody staining. Scale bars = 100 µm.

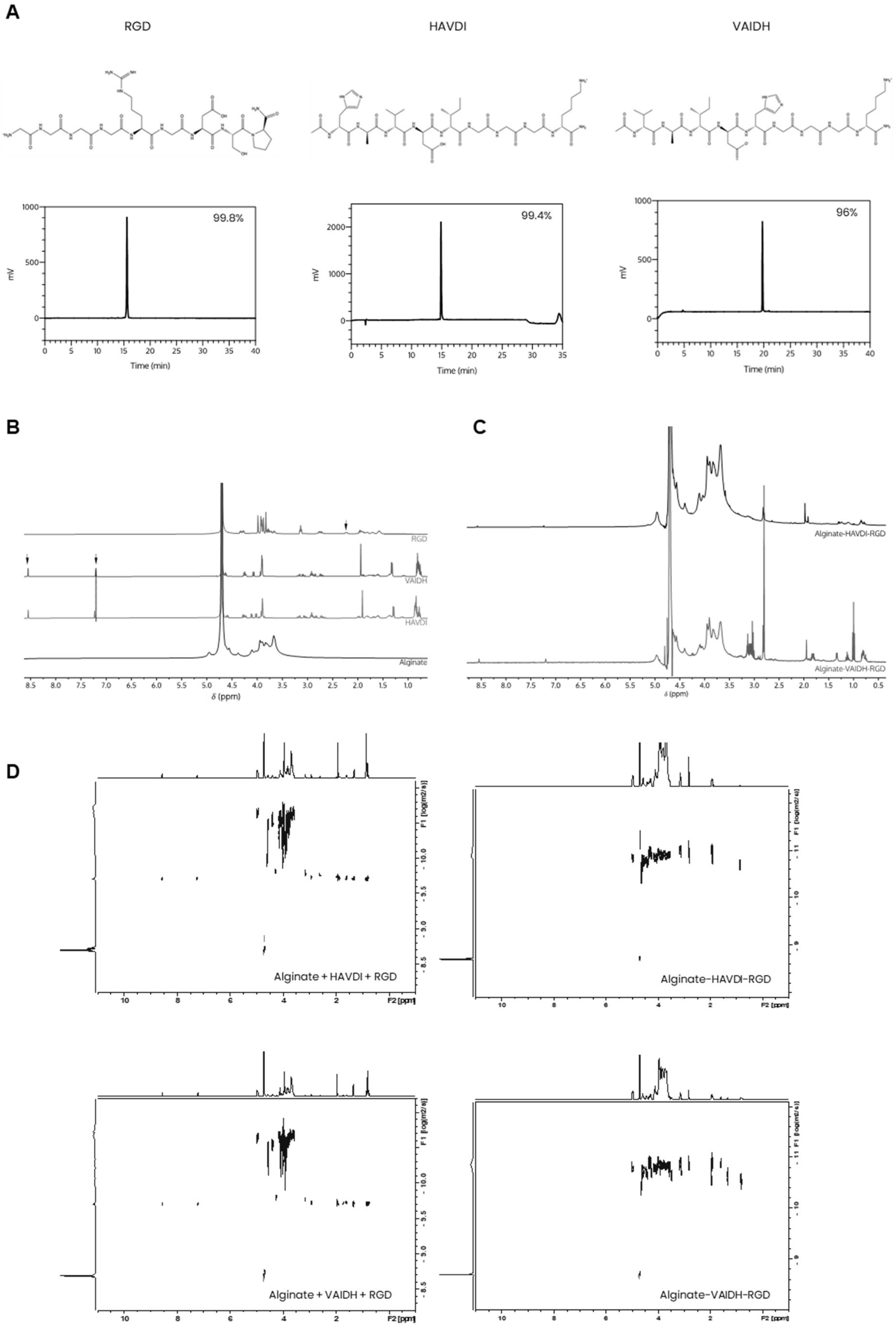
A
RGD
HAVDI
VAIDH
99.8%
99.4%
96%
mV
Time (min)
B
RGD
VAIDH
HAVDI
Alginate
δ (ppm)
C
Alginate-HAVDI-RGD
Alginate-VAIDH-RGD
δ (ppm)
D
Alginate + HAVDI + RGD
Alginate-HAVDI-RGD
Alginate + VAIDH + RGD
Alginate-VAIDH-RGD
F1 [log(m2/s)]
F2 [ppm]

**Figure S3 | Alginate modification: $^{1}$H NMR, DOSY $^{1}$H NMR and amplitude sweep analysis. A)** Structures and purity of peptides, determined by analytical HPLC. **B)** 1H-NMR spectra of individual RGD, VAIDH, HAVDI peptides, along with sodium alginate. Arrows indicate characteristic peaks of each peptide, while the dotted line highlights the shift in the imidazole ring peak between the scrambled (VAIDH) and HAVDI sequences. **C)** 1H-NMR spectra of alginate functionalized with HAVDI-RGD and VAIDH-RGD peptides. **D)** 1H-DOSY NMR of alginate modification, comparing a physical mixture of alginate and peptide powders (left) with covalently functionalized alginate following carbodiimide coupling (right). The physical mixture exhibits distinct diffusion coefficients for alginate and peptides, whereas the modified alginate shows a single diffusion coefficient, confirming covalent attachment of peptides to the polysaccharide chains.

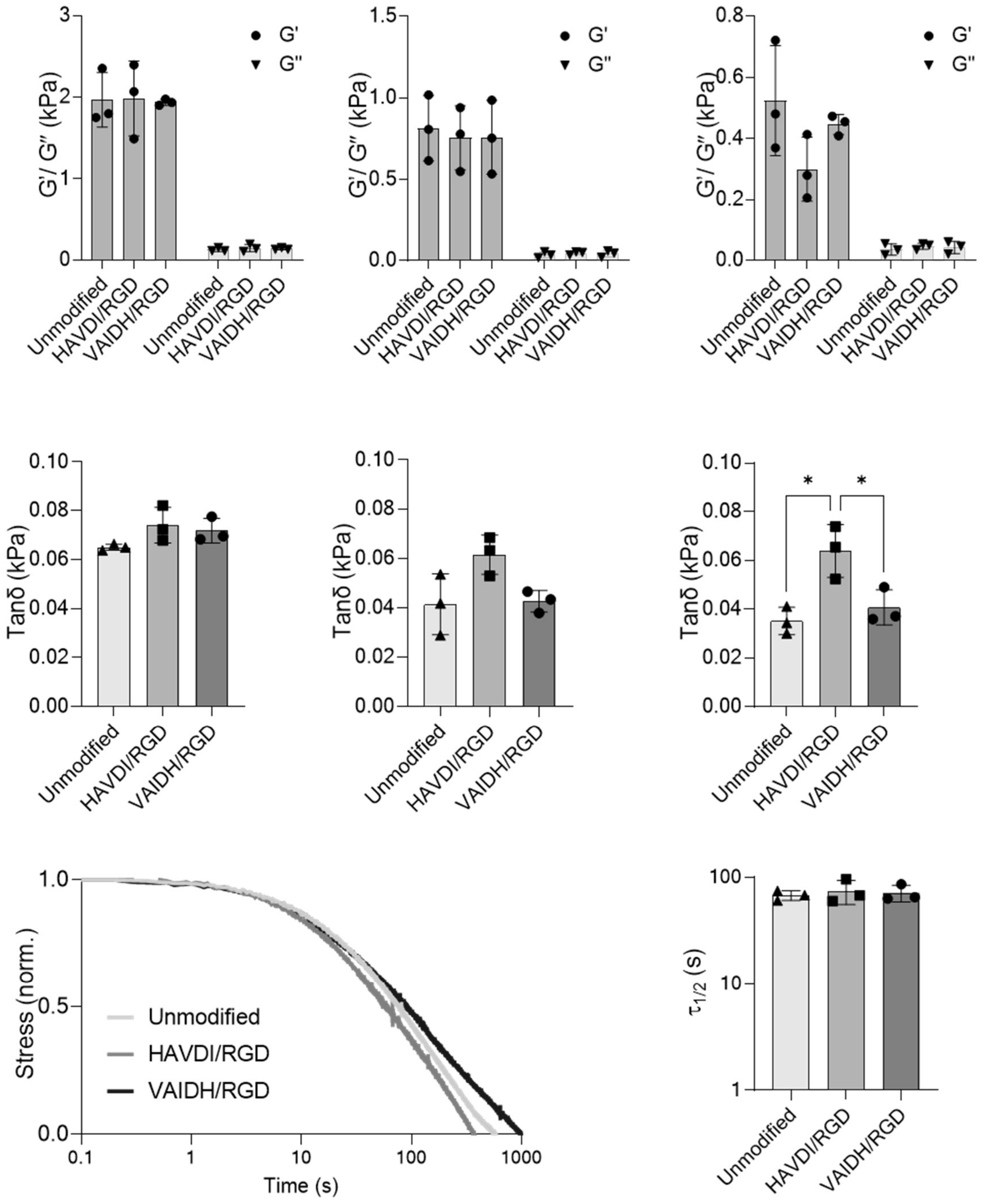


**Figure S4 | Rheological analysis of 2% (w/v) unmodified alginate and peptide-functionalized hydrogels in cell culture medium. A**) Stiffness measured as storage modulus (G') using amplitude sweep analysis, showing comparable mechanical behavior of hydrogels following peptide functionalization **B**) Tanδ measurements indicating that viscoelastic properties over time are influenced by peptide sequence. **C**) Stress relaxation analysis of alginate hydrogels after 24 hours in cell culture medium (15% strain; normalized data). D) Quantification of relaxation timescale from (C), showing the time required for gels to relax to half of their initial stress ($\tau_1/_2$). One-way ANOVA with Tukey's multiple comparisons post-hoc test was performed. Error bars represent mean ± s.d. (n = 3). Unmodified = 2% (w/v) unmodified alginate, RGD/HAVDI= 2% (w/v) alginate functionalized with RGD/HAVDI, RGD/VAIDH = 2% (w/v) alginate functionalized with RGD/VAIDH.
