## Supplementary Methods for "iPSC-Derived Chondroprogenitors as a Promising Cell Source for Cartilage Engineering: A Comparison with MSCs in Unmodified and Peptide-Functionalized Alginate Hydrogels"

### Peptide synthesis, $^1$H-NMR and $^1$H-DOSY Analysis

Three peptides were synthesised via microwave-assisted solid-phase synthesis (LibertyBlue, CEM Corporation, USA) at 0.5 mmol scale using standard Fmoc/tBu chemistry: Ac-HAVDIGGGK-$NH_2$ (HAVDI), its scrambled control Ac-VAIDHGGGK-$NH_2$ (VAIDH), and H-GGGGRGDSP-$NH_2$ (RGD). Synthesis employed L-amino acids coupled to Rink Amide MBHA resin (NovaBiotech). Unless noted, all chemicals were sourced from Sigma-Aldrich. Following peptide elongation on resin, the HAVDI peptide was acetylated by incubation with 200 µL N,N-diisopropylethylamine (DIPEA) and 100 µL acetic anhydride per 2 mL DMF for 20 minutes to acetylate the N-terminus, followed by three washes with DMF. For all peptides, the resin was washed with dichloromethane (DCM) prior to acidic cleavage using $H_2O$/triisopropylsilane (TIPS)/trifluoroacetic acid (TFA) (2.5:2.5:95) to remove protecting groups and release the peptides. Peptides were precipitated with chilled diethyl ether (~40 mL) for 10 minutes at -20 °C, centrifuged at 4,500 rpm for 10 minutes, dried under $N_2$ flow, and lyophilized. Crude peptides were dissolved (RGD and HAVDI: $H_2O$; VAIDH: 75:25 $H_2O$:MeCN) and purified by semi-preparative HPLC (Dionex P980 pump, UCV170U detector at 214/280 nm) on Phenomenex Gemini C18 (5 µm, 250 × 21.2 mm) at a flow rate of 8 mL/min). Purity was confirmed by reverse-phase HPLC using a Shimadzu system equipped with Phenomenex Luna C18(2) column (5 µm, 100 Å, 150 × 4.6 mm) at 1 mL/min. LC-MS analysis was performed using a Thermo Fisher Scientific LCQ Fleet quadrupole mass spectrometer with positive mode electrospray ionization (ESI+), equipped with Dr Maisch ReproSil Gold 120 C18 column (3 µm, 150 × 4 mm). Data were acquired using XCalibur 4.0 software. Purified peptides were lyophilized and stored at -20°C.

$^1$H-NMR spectroscopy was performed to characterise the structure of the coupled products, while diffusion-ordered spectroscopY ($^1$**H-**DOSY) was used to assess molecular diffusion of the and confirm the covalent conjugation. Both NMR and DOSY were performed using a Bruker Avance III 400 MHz spectrometer with a 5 mm BBO probe. Lyophilised functionalised alginate samples were dissolved at a concentration of 15 mg/mL in deuterium oxide ($D_2O$, 99.9 atom %, Sigma-Aldrich), sonicated, and transferred to NMR tubes. NMR spectra were recorded at 25 °C with 256 scans for $^1$H-NMR and 64 scans for $^1$**H-**DOSY. DOSY experiments were conducted using a Bruker sequence incorporating stimulated echo and bipolar gradient diffusion pulses. Gradient amplitudes ranged from 2% to 98% with a diffusion delay set to 1,000 µs. Data were processed using TopSpin 3.0 (Bruker BioSpin, USA) or MNova 15 (Mestrelab, USA).

### Supplementary Methods: Tables

| Component | Concentration/Amount | Supplier |
|---|---|---|
| **High-glucose DMEM (hgDMEM)** | Base medium | Thermo Fisher Scientific, UK |
| **Foetal Bovine Serum (FBS)** | 10% (v/v) | Thermo Fisher Scientific, UK |
| **Sodium Pyruvate (Na-pyruvate)** | 1% (v/v) | Thermo Fisher Scientific, UK |
| **MEM Non-Essential Amino Acids (MEM-NEAA)** | 1% (v/v) | Thermo Fisher Scientific, UK |
| **L-Glutamine** | 1% (v/v) | Thermo Fisher Scientific, UK |
| **Penicillin/Streptomycin (P/S)** | 1% (v/v) | Thermo Fisher Scientific, UK |
| **Amphotericin B** | 0.25 µg/mL | Thermo Fisher Scientific, UK |
| **Mesenchymal Stem Cell Growth Medium 2** | 10% (v/v) | PromoCell, Germany |

**Table SM1 | MSCs expansion medium composition.**

| Marker | Host species | Clone | Fluorochrome | Supplier | Cat. number |
|---|---|---|---|---|---|
| **CD45** | Mouse | HI30 | FITC | Biolegend | 982316 |
| **CD34** | Mouse | 581 | FITC | Biolegend | 343503 |
| **CD44** | Mouse | BJ18 | PE | Biolegend | 338807 |
| **CD90** | Mouse | 5E10 | Alexa Fluor 647 | Biolegend | 328116 |
| **CD146** | Mouse | P1H12 | BV421 | Biolegend | 361003 |
| **CD166** | Mouse | 3A6 | APC | Biolegend | 343905 |

**Table SM2 | Flow cytometry antibody panel.**

| Gene | Forward sequence | Reverse sequence |
|---|---|---|
| **ACTB** | 5’ – CGTGCTGCTGACCGAGG – 3’ | 5’ – GAAGGTCTCAAACATGATCTGGGT – 3’ |
| **SOX9** | 5’ – GCTCTGGAGACTTCTGAA – 3’ | 5’ – GGTACTTGTAATCCGGGTG – 3’ |
| **COL2A1** | 5’ – GGCTTCCATTTCAGCTATG – 3’ | 5’ – CAGTGGTAGGTGATGTTC – 3’ |
| **ACAN** | 5’ – GGCTTCCACCAGTGTGAC – 3’ | 5’ – GTGTCTCGGATGCCATACG – 3’ |
| **COL1A1** | 5’ – CAATGCTGCCCTTTCTGCTCC – 3’ | 5’ – CACTTGGGTGTTTGAGCATTGCCT– 3’ |
| **COL10A1** | 5’ – CGCTGAACGATACCAAATGC – 3’ | 5’ – CACTTGGGTGTTTGAGCATTG – 3’ |

**Table SM3 | QPCR primers used for human genes.**

| Primary Antibody | Concentration | Supplier | Cat. # |
|---|---|---|---|
| **Collagen type II** | 2 µg/mL | Thermo Fisher Scientific | MA512789 |
| **Aggrecan** | 1 µg/mL | Proteintech | 13880-1-AP |
| **Collagen type I** | 0.5 µg/mL | Proteintech | 67288-1-1G |

**Table SM4 | IHC primary antibodies.**

| **Symbol** | * | ** | *** | **** |
|---|---|---|---|---|
| **Meaning** | P ≤ 0.05 | P ≤ 0.01 | P ≤ 0.001 | P ≤ 0.0001 |

**Table SM5 | Statistical analyses symbols.**
